# Selection of the Early Genetic Code by Ultraviolet Light

**DOI:** 10.1101/2022.10.13.512048

**Authors:** Corinna L. Kufner, Stefan Krebs, Marlis Fischaleck, Julia Philippou-Massier, Helmut Blum, Dominik B. Bucher, Dieter Braun, Wolfgang Zinth, Christof B. Mast

## Abstract

The DNA sequences available in the prebiotic era were the genomic building blocks of the first life forms on Earth and have therefore been a matter of intense debate.^1,2^ On the surface of the Early Earth, ultraviolet (UV) light is a key energy source^3^, which is known to damage nucleic acids^4^. However, a systematic study of the sequence selectivity upon UV exposure under Early Earth conditions is still missing. In this work, we quantify the UV stability of all possible canonical DNA sequences and derive information on codon appearance under UV irradiation as selection pressure. We irradiate a model system of random 8mers at 266 nm and determine its UV stability via next-generation sequencing. As a result, we obtain the formation rates of the dominant dimer lesions as a function of their neighboring sequences and find a strong sequence selectivity. On the basis of our experimental results, we simulate the photodamage of short proto-genomes of 150 bases length by a Monte Carlo approach. Our results strongly argue for UV compatibility of early life and allow the ranking of codon evolutionary models with respect to their UV resistance.

## Introduction

Around 4.0 billion years ago^5^, short nucleic acids were exposed to UV irradiation as low as 200 nm on the surface of the Early Earth^6^ due to the absence of the atmospheric ozone layer. The absorption of UV photons can induce permanent lesions in short DNA strands, depending on the sequence, wavelength and the strand length^7^, which leads to different UV susceptibilities of each individual sequence. Before the evolution of complex repair enzymes^8^, photostability could have been a major survival criterion for the first oligonucleotides on the surface of the Early Earth which may have induced the selection of an initial sequence pool. Over time, less UV stable sequences could have evolved as enzymatic repair mechanisms emerged and became increasingly complex, targeted, and efficient. In this work, we determine the development of codon pairs under UV as selection pressure. Therefore, we measure the UV susceptibility of all possible DNA octamer (8mer) sequences experimentally by next-generation sequencing. The massively parallel approach allows us to monitor damaging interactions with respect to their sequence context. The 8mer model system mimics any short oligonucleotide which could have evolved on the surface of the Early Earth. Our work on the UV stability of short sequences extends the previous work on specific photolesion formation in modern genomic DNA^9,10^. We use our experimental data to simulate the UV susceptibility of a short proto-genome of 150 bases length (150mer) with a Monte Carlo approach. We simulate the stepwise development of codon pairs and determine an optimal codon chronology under UV exposure. This allows us to rank codon evolutionary models^2^ with respect to their UV resistance.

## Experimental Results

Our experimental approach is shown schematically in Figure 1a^12^: Random 8mers are irradiated at 266 nm and subsequently analyzed by next-generation sequencing. The polymerases used in our experiments stall at dimeric photolesions and damaged sequences are not amplified. A comparison to a non-irradiated sample allows us a background correction and to identify the photostability of each individual sequence. The photodegradation of example trimer (3mer) subsequences within the random 8mers is displayed by the dots in Figure 1b. By fitting the datapoints with a model (solid lines, equation system in SI) we are able to determine the damage formation of the 3mers, which range over several orders of magnitude depending on the sequence. Generally, sequences with a high dG content show a high photostability in contrast to dT-rich sequences. The biexponential decay of the d(TT) containing sequences indicates the formation of multiple photoproducts^13^. In the vicinity of d(TT), dG yields a higher photostability than dA, i.e. d(TTA) is less photostable than d(TTG), after absorbing 400 photons per base (P_p_B). On Early Earth^14^, an absorbed dose of around 35 P_p_B corresponds to the 250 nm – 280 nm photons absorbed by 10 μM mononucleotides^11^ underneath the surface (10 cm depth) of a low-absorption freshwater lake^3^ after 1 hour. The inlet in Figure 1b shows the depth profile of such an example scenario. Although the 3mer damage formation covers a wide range (Figure 1c), the majority of 3mers shows damage formation below 20·10^-3^ /P_p_B which is indicative of a high UV stability of most associated codons.

**Figure 1.**
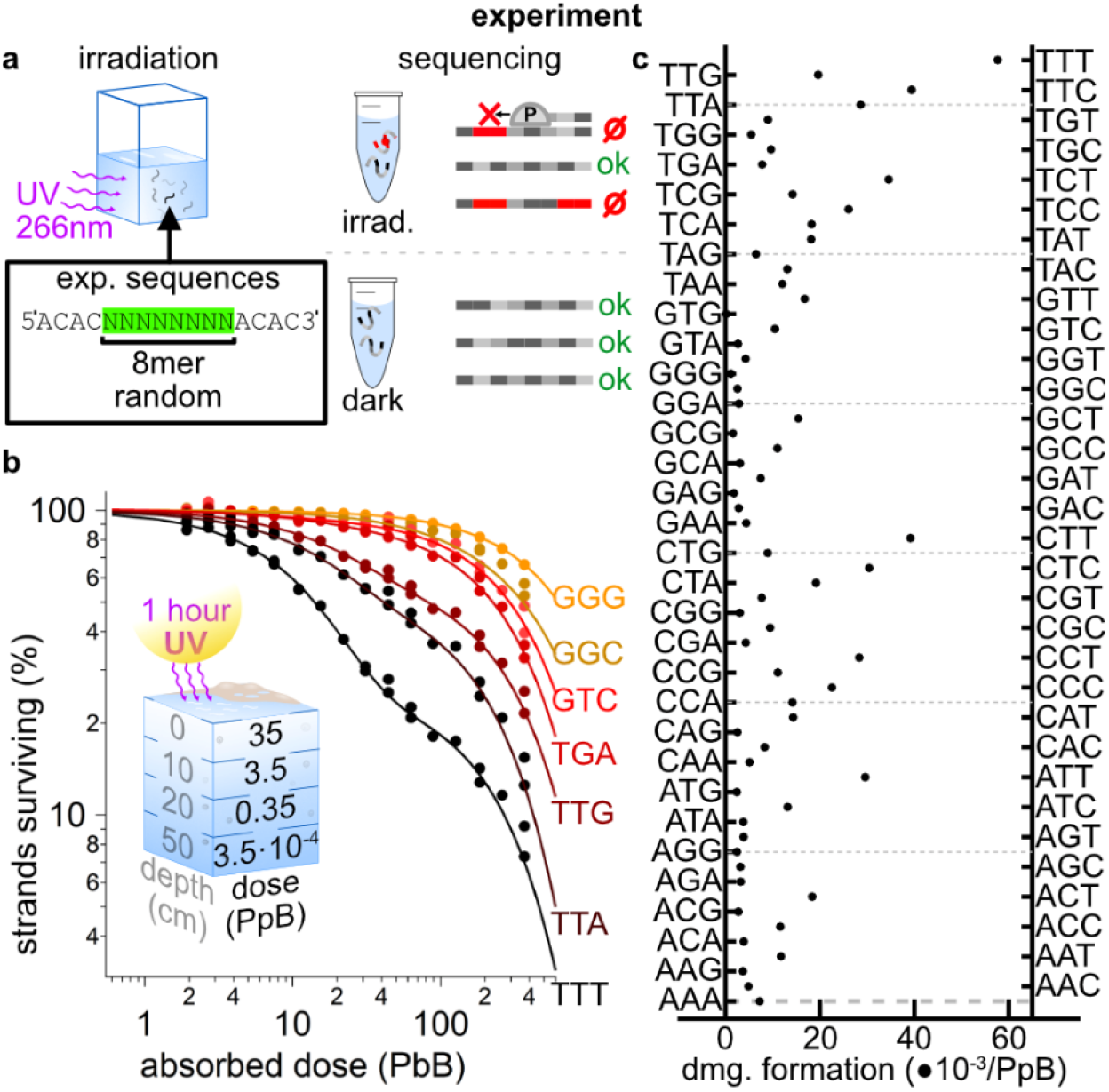
Experimental methods and results: (a) A solution of d(ACAC-NNNNNNNN-ACAC) sequences with 8 random canonical bases in the center (random 8mers) is irradiated at 266 nm. After 16 time intervals a small amount of the solution is removed and analyzed by next-generationsequencing. Damaged strands will not be detected in sequencing. (b) Percentage of surviving 3mers within the random 8mers as a function of the absorbed dose (dots) in units photons per base (P_p_B) and corresponding fit curves (lines). Inlet: Irradiation dose between 250 nm – 280 nm absorbed by 10 μM nucleotides^11^ at different depths of a prebiotic low-absorption freshwater lake^3^ after 1 hour (calculation in SI). (c) Damage formation of all possible sub-3mers within the random 8mers (axes left and right in alphabetical order).

## Simulation Results

For storage of genomic information and for the performance of functional tasks^15,16^, longer oligonucleotides need to be considered. However, due to the vast number of possible longer sequences, determining the UV stability of all sequences is experimentally not addressable. Based on our experimental 8mer data, we use a Monte Carlo approach to simulate the survival of single-stranded 150mer proto-genomes under UV exposure as little statistical differences are expected for strands longer than 150mers. Each proto-genome consists of 50 arbitrarily chosen codons. We model the development of an entire codon pool of 32 possible codon pairs (codon and anti-codon) under UV irradiation as sole selection pressure (Figure 2). Both codon and anti-codon are considered to emerge simultaneously as the complementary strand is temporarily required for replication and is therefore also exposed to UV light. We start by simulating the survival after 2 P_p_B of 10^4^ 150mers randomly (uniformly distributed) built from only one arbitrarily chosen codon pair (Figure 2, example table step 1). In each development step we add one codon pair to the pool of available codons from which we recalculate the averaged survival of all 150mer strands. We repeat this procedure until all 32 possible codon pairs are available to the pool (Figure 2, example table step 32). We call the sequence of available codon pools at each development step codon chronology. The choices of the starting codon pair (2 examples in Figure 3a) strongly affect the photostability of the codon pools within the entire codon chronology. 150mers consisting of the codon pair sequences d(TTT) and d(AAA) degrade on average almost 3 orders of magnitude faster under UV irradiation than 150mers consisting of d(GTG) and d(CAC). Examples for the surviving strands of the full 32 step codon chronologies at a dose of 2 P_p_B are shown in Figure 3b. The grey curves show randomly chosen codon chronologies and the blue curve represents a consensus chronology over various different development criteria from the literature^2^. The consensus chronology^2^ starts from the codon pair with the highest melting temperature of the corresponding amino acid. New codons are added at each development step, which can be obtained by a single mutation from the codon pool (Figure 2, grey letters in the table). The dashed curves are the most (green) and least (red) possible UV stable codon chronologies. At small codon pool sizes, there is a significant difference in the genomes surviving. All curves converge after 32 steps as the entire codon pool becomes available.

**Figure 2.**
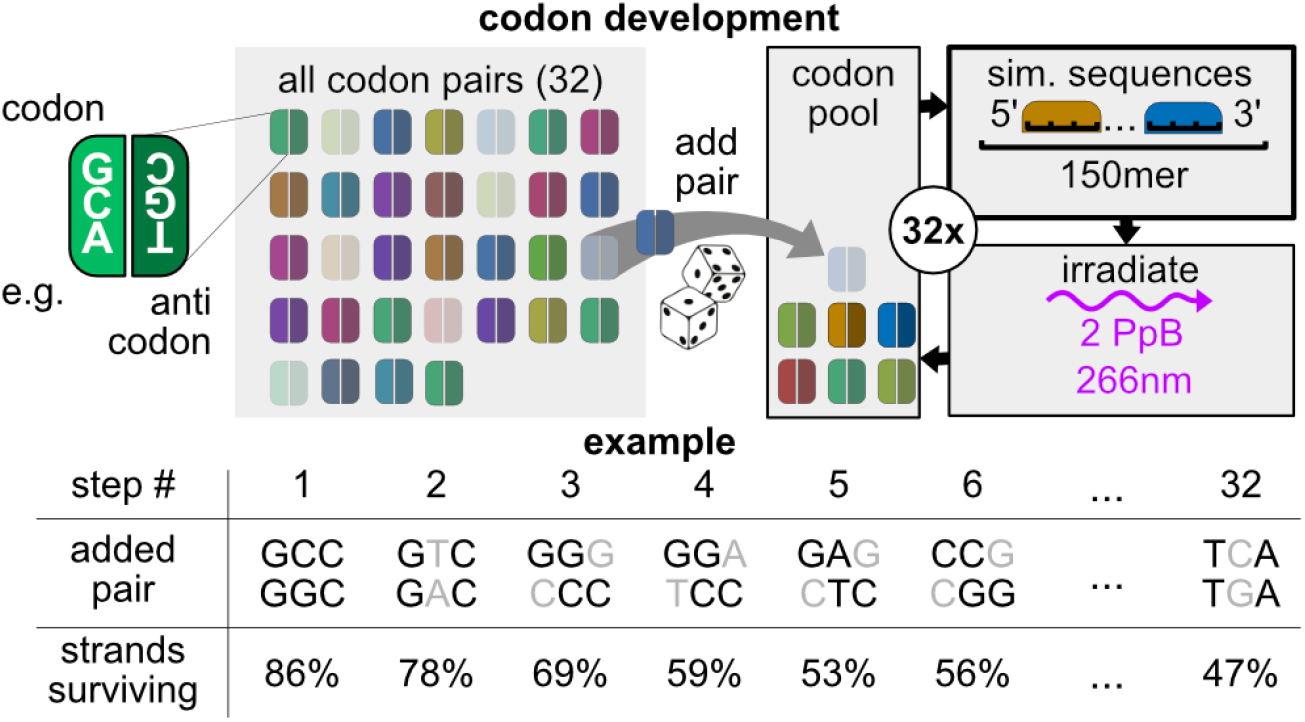
Simulation of the development steps of proto-genome (150mer): Codons and anti-codons are considered as pairs (32 total). A codon pool contains the codons available at a given time. In the first simulation step (see example table column 1), the codon pool contains only one statistically chosen codon pair. We simulate 10 000 strands of single-stranded 150mers arbitrarily consisting of the 3mer sequences of this codon pair and determine the averaged percentage of strands surviving an absorbed dose of 2 photons per base (P_p_B). We proceed by adding one available codon pair in each step and repeat the procedure until the codon pool consists of all 32 codon pairs. The example shows a codon chronology taken from^2^. Grey letters in the codon sequences that are added at each step indicate the change to sequences that are already available.

**Figure 3.**
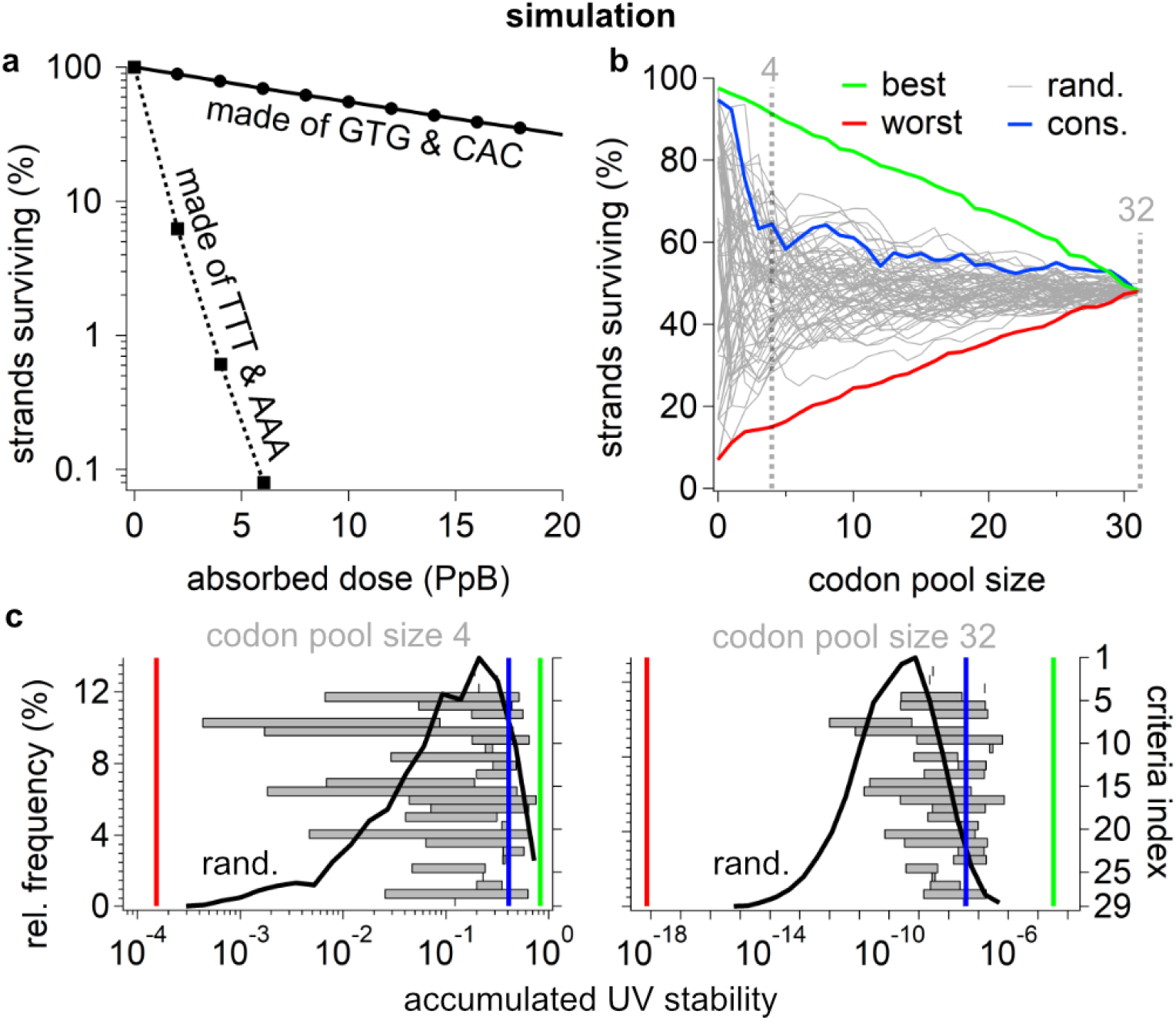
Results of the Monte Carlo simulation (a) Step 1: Percentage of strands surviving as a function of the absorbed dose in 2 averaged example proto-genomes (150mers) made of different starting codon pairs (dots: Monte-Carlo simulation, lines: exponential fit). (b) Full 32-step simulation (codon chronology): Percentage of strands surviving as a function of the codon pool size available to the proto-genome (150mer). Grey: statistical codon chronologies; green (most) and red (least) UV stable codon chronology found in this study; Blue: consensus chronology^2^. (c) Relative frequency histogram (left y-axes) of the accumulated UV stability (product of the strands surviving of all preceding steps of the corresponding codon chronology) of randomly chosen codon chronologies (black lines) at a codon pool size of 4 (left) and 32 (right). The accumulated UV stabilities of the most (green), least (red) UV stable and the consensus^2^ chronology (blue) are shown as vertical lines. The accumulated UV stability of codon chronologies which are associated with the literature criteria^1,2^ (right y-axes) are shown as grey horizontal bars. The detailed criteria index is shown in the SI.

Our simulated data can be used to estimate the accumulated UV stability of a codon chronology by multiplying the percentage of strands surviving cumulatively at each codon pool size (Figure 3c, x-axes). The accumulated UV stability quantifies the UV stability of the codon chronologies relative to each other. In a natural environment on the surface of the Early Earth, where codon pairs could have emerged randomly under UV exposure, the accumulated UV stability would have been the major survival criterion for the codon chronologies. The black lines in Figure 3c (left y-axis) show the relative frequency of the accumulated UV stability of a broad statistic of codon chronologies at an early (left: codon pool size of 4) and late (right: codon pool size of 32) development stage of the codon pool. The bounds of the UV stabilities of the most (green) and least (red) UV stable codon chronologies are shown as vertical lines for comparison. The amino acid criteria listed in the literature^1,2,17-20^ (right y-axis) are associated with multiple codon chronologies. Their accumulated UV stabilities range over several orders of magnitude (grey bars). The consensus chronology^2^ is shown as blue vertical line. At an early development stage (left), the accumulated UV stability of the literature criteria ranges from 10^-4^ to 10^-1^, which is within the range of the random chronologies. At late stages (right) the accumulated UV stability of random codon chronologies ranges from 10^-20^ to 10^-6^ whereas the literature criteria^1,2^ range from 10^-13^ to 10^-7^. This means that the codon chronologies following the literature criteria^2^, have high accumulated UV stabilities despite being unrelated to UV light as selection pressure. Remarkably, this is reflected in the high accumulated UV stability of the consensus chronology^2^ not only at late development stages (32 steps), but already at early stages (4 steps). Despite the absence of UV exposure as an explicit selection criterion, the consensus chronology^2^ starts with very photostable codon pairs (Figure 3b).

Table 1 shows the most UV stable codon chronology (green line in Figure 3) as an example. In each line, the listed codon pair (column 2) is added to the codon pool and yields a percentage of strands surviving (column 3). The rank boxes represent groups of codon pairs with very similar survival rates. Within each rank box (column 1) codon pairs can be permutated and will yield the same percentage of strands surviving (column 3). Guanine-rich codon pairs can be found at the low rank numbers with high UV stability and di-pyrimidine-rich codon pairs at high rank numbers with low UV stability. At low ranks, single T and CC sequences are flanked by G.

**Table 1.**
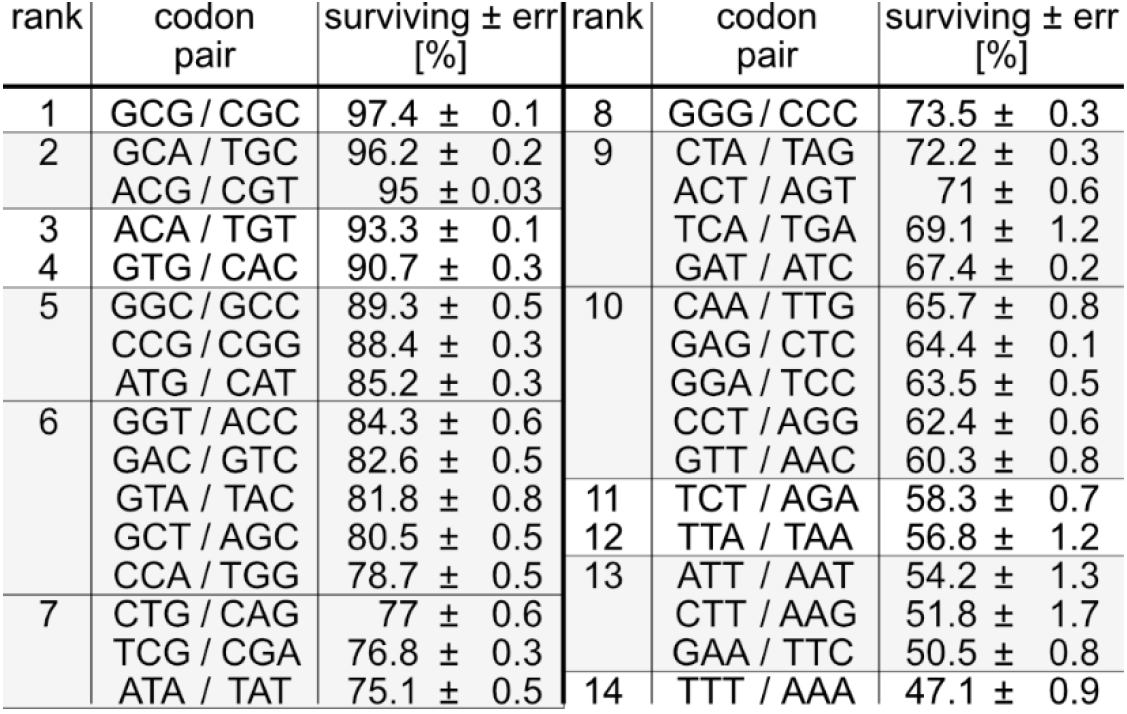
Example of the most UV-stable codon chronology (green lines in Figure 3). The codon pairs within each rank box group have very similar survival rates and can therefore be permutated without changing the percentage of strands surviving in the codon chronology.

## Discussion

The composition of the genome pools available to the first forms of life on Early Earth is a challenge for the models of the first organisms. Particularly, environmental impacts could have influenced the survival of early sequences. Ultraviolet light is an important energy source on the surface of the Early Earth and could have imposed a selection pressure on oligonucleotides.

With UV stability as sole selection pressure and without further biases, we study the survival of all possible hexamers on a nearest and next-to-next neighbor interaction level, mimicking the selection from a random sequence pool on Early Earth. We find Guanine-rich sequences could have dominated the sequence pool available to early genomes. The high UV stability of Guanine plays an important role for the selectivity. Guanine-containing sequences have been shown to promote sequence selective self-repair^21,22^ of pyrimidine dimer lesion which could have been a dominant repair pathway prior to the evolution of enzymatic repair. In the vicinity of pyrimidines^23^, the efficient formation of charge separated states with ultrashort lifetimes on the order of several 10 picoseconds could have reduced the formation of adjacent pyrimidine dimer lesions and thereby exhibited a protective function. However, the G content alone is not sufficient to determine the impact of UV radiation. Instead, the full sequence context has to be considered. We perform this by simulating the survival of 150mer proto-genomes and find an optimal codon chronology under UV exposure. Interestingly, our optimal codon chronology is widely compatible with other selection criteria reported in the literature ^1,2,17-20^ and the consensus chronology^2^. Under exposure to UV light, these findings increase the probability of the consensus chronology^2^ with an independent selection criterion. Our system can be transferred to extraterrestrial environments and has the potential to predict the development of a codon pool on exoplanets under exposure to UV irradiation. In a meteoritic setting^24^, UV light could even have been the dominant selection criterion. Our major findings support the big idea that if oligonucleotides have developed on the surface of the Early Earth, UV-induced selection processes have likely played a role from early stages onward.

## Materials and Methods

The DNA 5’-ACAC-NNNNNNNN-ACAC-3’ sequences (random 8mers) were purchased from biomers.net (Germany) and dissolved (1 mM nucleobase concentration, optically thick) in PBS buffer (1x). 3.45 mL of the solution were irradiated at 266 nm by a nanosecond pulsed laser (AOT-YVO-25QSP/NOPA, AOT, UK, repetition rate 6.5 kHz, average power 20 mW) in a 10 mm quartz cuvette (type 117100F-10-40, Hellma, Germany). During irradiation the solution was continuously recirculated. After irradiation of the desired dosages, we removed 50 μL of the solution 16 times, which corresponds to 16 different doses between 2 and 560 photons per base (P_p_B). We prepared the collected samples for next-generation-sequencing with a commercial library preparation kit (Swift Accel-NGS 1S DNA library kit, Swift Bioscience, USA) following the protocol. We measure the reduction in read frequency for photodamaged strands vs. intact random 8mers^12^.

## Supporting information

Monte Carlo Simulation (Labview)

## Data Availability

The data that support the findings of this study are available within the paper an its Supplementary Information.

## Author Contributions

C. B. M., W. Z., D. B., D. B. B. and C. L. K. conceived the project and directed the research. C. B. M. prepared the samples. C. L. K. and W. Z. conducted the irradiation experiments. S. K., M. F., J. P.-M. and H. B. performed the next-generation sequencing. C. B. M. and W. Z. analyzed the data. C. L. K., C. B. M, and D. B. explored prebiotic implications. C. B. M., W. Z., D. B., D. B. B., S. K. and C. L. K. wrote the paper.

## Acknowledgements

The authors are grateful to J. D. Sutherland for answers to questions and discussions related to the topic. The authors further thank G. G. Lozano for comments on the draft of this article.

## Funding Information

This work was supported by the Deutsche Forschungsgemeinschaft (DFG, German Research Foundation) through the Clusters of Excellence “Center of Integrated Protein Science Munich” (W.Z.) and Project-ID 364653263 – TRR 235 (CRC235), Project P08 (C.B.M). Funding from the Simons Foundation (327125 to D.B., SCOL 290360FY18 to Dimitar Sasselov), Volkswagen Initiative ‘Life? – A Fresh Scientific Approach to the Basic Principles of Life’ (C.B.M., D.B.), ERC ADV 2018 Grant 834225 (EAVESDROP) (D.B.) and from ERC-2017-ADG from the European Research Council (D.B.) is gratefully acknowledged. Funded by the Deutsche Forschungsgemeinschaft (DFG, German Research Foundation) under Germany’s Excellence Strategy – EXC-2094 – 390783311 (D.B., C.B.M). The work is supported by the Center for Nanoscience Munich (CeNS).

## Competing Interests

No competing interests exist.

## Additional Information

## Supplementary Information

**Correspondence and requests for material** should be addressed to Christof B. Mast.

## Notes

### Competing Interest Statement

The authors have declared no competing interest.

## References

1 Trifonov, E. N. Consensus temporal order of amino acids and evolution of the triplet code. Gene 261, 139–151 (2000).

2 Trifonov, E. N. The triplet code from first principles. Journal of Biomolecular structure and dynamics 22, 1–11 (2004).

3 Ranjan, S. et al. UV Transmission in Natural Waters on Prebiotic Earth. Astrobiology 22, 242–262 (2022).

4 Downes, A. & Blunt, T. P. Proceedings of the Royal Society London 26, 488–500 (1877).

5 Joyce, G. F. The antiquity of RNA-based evolution. Nature 418, 214–221 (2002).

6 Ranjan, S., Wordsworth, R. & Sasselov, D. D. The surface UV environment on planets orbiting M dwarfs: implications for prebiotic chemistry and the need for experimental follow-up. The Astrophysical Journal 843, 110 (2017).

7 Görner, H. New trends in photobiology: Photochemistry of DNA and related biomolecules: Quantum yields and consequences of photoionization. Journal of Photochemistry and Photobiology B: Biology 26, 117–139 (1994).

8 Sancar, A. Mechanisms of DNA repair by photolyase and excision nuclease (Nobel Lecture). Angewandte Chemie International Edition 55, 8502–8527 (2016).

9 Hu, J., Adar, S., Selby, C. P., Lieb, J. D. & Sancar, A. Genome-wide analysis of human global and transcription-coupled excision repair of UV damage at single-nucleotide resolution. Genes & development 29, 948–960 (2015).

10 Lu, C., Gutierrez-Bayona, N. E. & Taylor, J.-S. The effect of flanking bases on direct and triplet sensitized cyclobutane pyrimidine dimer formation in DNA depends on the dipyrimidine, wavelength and the photosensitizer. Nucleic Acids Research 49, 4266–4280 (2021).

11 Pearce, B. K., Pudritz, R. E., Semenov, D. A. & Henning, T. K. Origin of the RNA world: The fate of nucleobases in warm little ponds. Proceedings of the National Academy of Sciences 114, 11327–11332 (2017).

12 Kufner, C. L. et al. Sequence Dependent UV Damage of Complete Pools of Oligonucleotides. bioRxiv, 2022.2008.2001.502267 (2022).

13 Lemaire, D. G. & Ruzsicska, B. P. Quantum yields and secondary photoreactions of the photoproducts of dTpdT, dTpdC and dTpdU. Photochemistry and photobiology 57, 757–769 (1993).

14 Ranjan, S. & Sasselov, D. D. Constraints on the early terrestrial surface UV environment relevant to prebiotic chemistry. Astrobiology 17, 169–204 (2017).

15 Longo, L. M. et al. Primordial emergence of a nucleic acid-binding protein via phase separation and statistical ornithine-to-arginine conversion. Proceedings of the National Academy of Sciences 117, 15731–15739 (2020).

16 Su, M., Ling, Y., Yu, J., Wu, J. & Xiao, J. Small proteins: untapped area of potential biological importance. Frontiers in genetics 4, 286 (2013).

17 Li, D. J. Formation of the Codon Degeneracy during Interdependent Development between Metabolism and Replication. Genes 12, 2023 (2021).

18 Ikehara, K., Omori, Y., Arai, R. & Hirose, A. A novel theory on the origin of the genetic code: a GNC-SNS hypothesis. Journal of molecular evolution 54, 530–538 (2002).

19 Brooks, D. J. & Fresco, J. R. Increased frequency of cysteine, tyrosine, and phenylalanine residues since the last universal ancestor. Molecular & Cellular Proteomics 1, 125–131 (2002).

20 Eigen, M. & Schuster, P. The hypercycle: a principle of natural self-organization. (Springer Science & Business Media, 2012).

21 Bucher, D. B., Kufner, C. L., Schlueter, A., Carell, T. & Zinth, W. UV-Induced Charge Transfer States in DNA Promote Sequence Selective Self-Repair. J. Am. Chem. Soc. 138, 186–190, doi:10.1021/jacs.5b09753 (2016).

22 Pan, Z., Chen, J., Schreier, W. J., Kohler, B. & Lewis, F. D. Thymine dimer photoreversal in purine-containing trinucleotides. The Journal of Physical Chemistry B 116, 698–704 (2011).

23 Kufner, C. L., Zinth, W. & Bucher, D. B. UV-Induced Charge-Transfer States in Short Guanosine-Containing DNA Oligonucleotides. ChemBioChem 21, 1–6 (2020).

24 Kvenvolden, K. A., Lawless, J. G. & Ponnamperuma, C. Nonprotein amino acids in the Murchison meteorite. Proceedings of the National Academy of Sciences 68, 486–490 (1971).

