## Supplementary methods and text for "Selection of the Early Genetic Code by Ultraviolet Light"

### Supplementary Information

#### Table of contents:

**Experiment:**

SI-1: Materials and methods for illumination and sequencing of damaged DNA strands

SI-2: Analysis of sequencing data

SI-3: Approximation of absorbed dose

SI-4: Dimeric and trimeric damage rates

SI-5: Context-dependent attenuation of the damage rates

**Monte Carlo simulation:**

SI-6: Numerical damage model on random DNA pools

SI-7: Evolutionary algorithm for calculation of codon chronologies with worst and optimal UV resistance

SI-8: Deduction of codon chronologies from amino acid chronologies

SI-9: Strands surviving for reference codon chronologies

References

#### Experiment

##### SI-1: Materials and methods for illumination and sequencing of damaged DNA strands

To model UV damage of long strands of arbitrary sequence, it was necessary to first measure the formation of the most abundant dimeric photolesions per adsorbed photon (damage rate) under well-defined experimental conditions for all permutations of neighboring bases. While the formation of dimeric lesions has been studied in detail under various different conditions (e. g. cyclobutane pyrimidine dimers as a function of the adjacent bases)[1-5], a uniform set of dimeric damage rates including adenine-adenine photoproducts as a function of all neighboring bases under uniform conditions, has not been available so far.

However, the determination of the individual damage rates for each of the dominant dimer lesions of two pyrimidine bases (TT, CT, TC, CC) or two adenosine bases (AA), and for each possible neighbor sequence would require $4^4$ (2 neighbors on each side) x 10 (dimer combinations without order) x 12 (typically required single dose values) = 30720 singular irradiation experiments and downstream HPLC runs. To avoid this, we chose instead the irradiation of single pools of randomized, short DNA strands with subsequent analysis through high throughput sequencing.

We prepared the samples from lyophilized commercially synthesized single stranded DNA (ssDNA) with a length of 16 nucleotides and sequence `5′-ACACNNNNNNNNACAC-3′` (biomers, Germany) by dissolution in spectroscopically pure water (LC-MS grade, Carl-Roth) and titration to 1x PBS buffer conditions (137 mM NaCl, 2.7 mM KCl, 10mM $Na_2HPO_4$, 1.8 mM $KH_2PO_4$) to maintain a pH of 7.4 and a DNA base concentration of 1 mM. The samples were centrifuged, vortexed and the final concentration was checked by Nanodrop (Thermofischer, US) and UV-Vis measurement (Shimadzu UV1800). The DNA sequences consist of three sequence parts: a central part containing 8 random canonical nucleotides and two flanking parts of the sequence d(ACAC), which act as recognition tags in sequencing analysis.

3.45 ml of buffered DNA solution were irradiated in a fused silica cuvette (type 117100F-10-40, Hellma, Germany) with a path length of 10 mm. During irradiation, the cuvette was kept at a constant temperature of 22°C and mixed by magnetic stirring. The sample was irradiated by a Nd-based laser system (AOT-YVO-25QSP/NOPA, AOT, UK) with a repetition rate of 6.5 kHz, an average power of 20 mW, at a wavelength of 266 nm. The laser power was checked before and after passing through the sample using beam splitters and appropriately placed power meters (Ophir, Israel). The difference between the two readings was used to determine the total absorbed dose in each case. After the desired dose was reached, exposure was interrupted for a short time to collect 50 µl of sample, which was then frozen at -80 °C for subsequent high-throughput sequencing. 12 samples in total were collected in duplicate with exponentially increasing doses from 2 to 560 photons per base. Residual non-irradiated DNA solution was kept as control.

All samples were prepared for sequencing according to the standard protocol of the Swift Accel-NGS 1S DNA library kit (Swift Bioscience, USA). Special care was taken to process the samples immediately after thawing to avoid thermal alteration of the damage state of the strands. Sequencing was performed using a Hi-Seq high-throughput sequencer (Illumina, USA) with a planned number of 40 to 60 million reads per sample.

**SI-2: Analysis of sequencing data**

The principle for determining context-dependent damage rates of sequences is based on the inherent property of the high-fidelity polymerases used for sequence library generation to stall the polymerization process when noncanonical bases are present[6]. Using a complete random sequence pool (every possible sequence of the central randomized part of the DNA occurs at least once) leads to a detectable decrease in the amount of successful sequencing reads for sequences containing UV lesions. Since we have assigned the amount of sequencing reads to be 40 to 60 million and $4^8 = 65536$ different sequences are possible for a sequence with 8-random nucleotides, each possible sequence will ideally occur about 1000 times in our unexposed sample. This means that the dynamic range for UV damage detection will span approximately 3 orders of magnitude, enabling us to identify the most common UV lesions that occur more frequently than $10^{-3}$ occurrences per strand in a complete sequence context. Due to sequence bias in synthesis and sequencing, we expect significant variations for each sequence, which are, however, identical for all samples used since they were obtained from the same stock solution.

Thus, the only difference between the unexposed control and the exposed samples is due to the reduction in the amount of reads due to the stalling effect of the polymerase used on the damaged sequences. The inherent sequence bias from synthesis and sequencing can therefore be eliminated by normalizing the amount of reads $Q_i(D)$ per 16-mer sequence $i$ for a sample that was exposed to the dose D with the amount of reads $Q_i(0)$ of the non-exposed control sample to obtain the survival probability $S(i, D) = Q_i(D)/Q_i(0)$. In the simplest case, such as for low exposure doses, one can assume that the decrease in the amount of undamaged strands $dQ_i$ is proportional to their current number $Q_i$, the differential dose $dD$ and the global damage rate $\mu_i$ which translates to the linear differential equation:

(SI-e 1) $$\frac{dQ_i}{dD} = -\mu_i \cdot Q_i$$

With the monoexponential solution:

(SI-e 2) $$Q_i(D) = Q_i(0) \cdot \exp(-\mu_i \cdot D)$$

For higher dosages, other damage states and back-reactions need to be taken into account especially if the sequence $i$ contains a larger number di-pyrimidines. In this case, a bi-exponential model applies (see Figure 1b for e.g. TTT trimers) [7]. In the calculations below, the former case with low dose values in the range of 2 photons per base applies, which is why it is sufficient to use the initial slope of the simple exponential model SI-e2 as the damage rate.

Since we are not only interested in the damage rates of the entire strand but also in all subsequences, $i$ can also be chosen as a shorter sequence in a central position. In this case, the number of undamaged strands containing a central subsequence $i$ after a dose $Q_i(D)$ is then obtained simply by summing up all sequence reads that contain the subsequence i at the center position. In addition, the part of the damage that is not formed within sequence i must then be excluded. We achieve this by a normalization step with respect to a non-damaging sequence $i_G$=poly-G of the same length as i.

(SI-e 3) $$S(i,D) = \left[\frac{Q_i(D)}{Q_i(0)}\right] / \left[\frac{Q_{i_G}(D)}{Q_{i_G}(0)}\right],$$

$Q_{i_G}(D)/Q_{i_G}(0)$ is the number of surviving strands determined from sequencing that have a poly-G sequence at the central position that is known to be negligibly damaged at doses up to 500 photons per base [7]. The corresponding data for subsequences of length 3 (trimers) is shown in the main text Figure 2b (dots) including the bi-exponential fits (lines).

Irrespective of $i$ being a subsequence or the entire sequence of the strand, the molecular damage rates of the respective dimer damage can now be calculated from the damage rates. For this purpose, we split the damage rates using the example of the monoexponential damage model SI-e 2, here for a trimer subsequence $i = XYZ$ with XYZ being a specific sequence made from canonical bases:

(SI-e 4) $$\mu_i = \Delta\mu_{\mathrm{X}} + \Phi_{XY} + \Phi_{YZ} + \Delta\mu_Z$$

where $\Delta\mu_X$ and $\Delta\mu_Z$ denote contributions that arise from possible lesion formations with the adjacent edge sequences. If we would consider only isolated trimer strands and not 16mere as here in the experiment, these contributions would lead to an effective increase of the damage rates, since absorbed photons can only lead to damage formation within the strand $\Delta\mu_X = 0.5 \cdot \Phi_{XY},\ \Delta\mu_Z = 0.5 \cdot \Phi_{YZ}$. In our case of considering of sub-sequences, e.g. 3 within the 16mers strands, the edge interaction results from all possible molecular damage possibilities with the nearest adjacent base:

(SI-e 5) $$\Delta\mu_X = 0.25 \cdot (\Phi_{AX} + \Phi_{CX} + \Phi_{GX} + \Phi_{\mathrm{TX}}),$$

$$\Delta\mu_Z = 0.25 \cdot (\Phi_{ZA} + \Phi_{ZC} + \Phi_{ZG} + \Phi_{\mathrm{ZT}})$$

The global damage rates $\mu_i$ are now determined by the fit function SI-e 2. This procedure is valid for the low-dose case used in this work (2 PpB) allowing for virtually independent damage event. For all possible sequences $i = XYZ$, SI-e 4 defines a system of linear equation which can be written in general terms as

(SI-e 6) $$\mu_i = \sum_{ij} A_{ij} \cdot \Phi_j$$

with $j$ being all possible dimer sequences. The solution of this equation system yields the molecular dimeric damage rates $\Phi_j$ presented in the next chapter. To determine the error of the molecular damage rates $\Delta\Phi_j$, the SI-e 6 system was solved for 1000 different $\mu_i'$ values, which lie within the fit-error around the fitted rate $\mu_i$. The width of the resulting Gaussian distribution of solutions $\Phi_j'$ then defines the error $\Delta\Phi_j$.

#### SI-3: Approximation of adsorbed dose

In order to determine the number of actually absorbed photons $N_{abs}$, the total number of bases $N_{base}$ and thus the real dose $D = N_{abs}/N_{base}$ of the respective samples, the power difference $W_{abs}$ of the laser beam in front of and behind the sample is determined over the entire measurement period. Here, reflections at the cuvette are taken into account by prior calibration. The number of absorbed photons is thus $N_{abs} = \int W_{abs}\, dt/h\nu$.

The actual number of bases is determined from the strand concentration of the 16mers after sample preparation by a dedicated UV absorbance measurement (see SI-1). Due to the designed sequence bias caused by the alternating edge sequences, the extinction coefficient $\epsilon_{16}$ is determined by piecewise averaging over all pairwise base combinations[8] $\epsilon_{i,j}, i,j \in \{A,C,G,T\}$:

(SI-e 7) $$\epsilon_{16} = \frac{\sum_{i=1}^{15}\varepsilon_{i,i+1} - \sum_{i=2}^{15}\varepsilon_{i}}{16} = \frac{1}{16}\cdot\left(\epsilon_{A,C} + \epsilon_{C,A} + \epsilon_{A,C} + \epsilon_{C,N} + 7\epsilon_{N,N} + \epsilon_{N,A} + \epsilon_{A,C} + \epsilon_{C,A} + \epsilon_{A,C} - 3\epsilon_{A} - 3\epsilon_{C} - 8\epsilon_{N}\right)$$

With $\epsilon_N = \frac{\varepsilon_A+\varepsilon_C+\varepsilon_G+\varepsilon_T}{4}$, $\varepsilon_{C,N} = \frac{\varepsilon_{C,A}+\varepsilon_{C,C}+\varepsilon_{C,G}+\varepsilon_{C,T}}{4}$, $\varepsilon_{N,A} = \frac{\varepsilon_{A,A}+\varepsilon_{C,A}+\varepsilon_{G,A}+\varepsilon_{T,A}}{4}$ and $\varepsilon_{N,N} = 1/16\cdot\sum\epsilon_{i,j}$.

The total number of bases $N_{base}$ is then calculated using the Lambert-Beer Law: $N_{base} = V\cdot N_A\cdot\frac{A_{260}}{\epsilon_{16}\cdot d}$ from the absorbance $A_{260}$ at the wavelength $\lambda = 260nm$ in the sample cell with a path length $d$ in a separate experiment.

**Table SI-t 1.** Calculated number of effective doses per sample

| Sample # | 1 | 2 | 3 | 4 | 5 | 6 | 7 | 8 | 9 | 10 | 11 | 12 |
|---|---|---|---|---|---|---|---|---|---|---|---|---|
| Dose (photons/base) | 2 | 15 | 22 | 31 | 45 | 63 | 92 | 127 | 184 | 261 | 370 | 559 |

*Approximation of absorbed dose in an early life context*

To approximate the magnitude of absorbed UV radiation in the context of an early Earth, we use a warm-little-pond model. We assume a maximum base mass-fraction of about *1 ppm*, $c{\sim}10\ \mu M$ in water, leached from a carbonaceous meteroid[9] to obtain the depth dependence shown in Figure 1b. Using Lambert-Beer's law, we deduce the total number of photon absorptions per time:

(SI-e 8) $$N = \int I(\nu)\frac{1}{h\nu}\,\epsilon(\nu)\cdot\ln(10)\cdot\frac{1}{N_A\cdot 10^{-3}}d\nu$$

with the intensity spectrum $I(\nu)$ of an early Earth[10] (zenith angle: 0°, albedo: ocean), the photon frequency $\nu$, the Planck constant $h$, the extinction coefficient $\epsilon(\nu)$ of thymine[11], and the Avogadro constant $N_A$. We perform the integral between 250 and 280 nm and yield $N = 35\,photons/hour$.

#### SI-4: Dimeric and trimeric damage rates

Using the procedure described in section SI-2, we were able to determine the molecular damage rates $\Phi_i$ for all dimers $i$ and damage rates $\mu_i$ for all possible trimers $i$. The detection of the specific damage type was of secondary importance here, since it is assumed that the dimer damage considered here, can at least affect the readout processes in a proto-genomes due to its property to impair the function of modern polymerases. Thus, we obtain the following values for the effective damage rates of the dimers from solving SI-e 6.

**Table SI-t2:** Molecular damage rates $\Phi_{i,j}$ for dimers:

| Sequence i | TT | TC | CC | AA | AT | AC | GC | AG | GG | GT |
|---|---|---|---|---|---|---|---|---|---|---|
| Rate $\Phi_i$ ($10^{-3}$ dmg/photon) | 20 | 10 | 5 | 2 | $< 10^{-3}$ | $< 10^{-3}$ | $< 10^{-3}$ | $< 10^{-3}$ | $< 10^{-3}$ | $< 10^{-3}$ |
| Error ($10^{-3}$ dmg/photon) | 2 | 1 | 0.5 | 0.2 | 0.5 | 0.5 | 0.4 | 0.6 | 0.3 | 0.1 |

The normalized damage rates $\mu_i$ of the trimers shown in main text Figure 2c are determined using the fitting procedure with the mono model SI-e 2, shown in table SI-t3.

**Table SI-t3:** Damage rates $\mu_i$ for all possible trimer sequences (cf. Figure 2c).

| 3-mer | $\mu_i$ ($10^{-3}$dmg/photon) | 3-mer | $\mu_i$ ($10^{-3}$dmg/photon) | 3-mer | $\mu_i$ ($10^{-3}$dmg/photon) | 3-mer | $\mu_i$ ($10^{-3}$dmg/photon) |
|---|---|---|---|---|---|---|---|
| AAA | 7.2 | TAA | 12.0 | GAA | 4.4 | CAA | 5.1 |
| AAC | 4.8 | TAC | 13.1 | GAC | 2.8 | CAC | 8.3 |
| AAG | 3.7 | TAG | 6.5 | GAG | 1.8 | CAG | 2.5 |
| AAT | 11.8 | TAT | 18.1 | GAT | 7.5 | CAT | 14.3 |
| ACA | 3.9 | TCA | 18.2 | GCA | 3.1 | CCA | 14.1 |
| ACC | 11.6 | TCC | 26.1 | GCC | 11.0 | CCC | 22.5 |
| ACG | 2.8 | TCG | 14.2 | GCG | 1.6 | CCG | 11.1 |
| ACT | 18.3 | TCT | 34.5 | GCT | 15.4 | CCT | 28.3 |
| AGA | 3.2 | TGA | 7.7 | GGA | 2.9 | CGA | 4.3 |
| AGC | 3.1 | TGC | 9.6 | GGC | 2.5 | CGC | 9.4 |
| AGG | 2.4 | TGG | 5.4 | GGG | 1.1 | CGG | 3.0 |
| AGT | 3.8 | TGT | 9.0 | GGT | 4.2 | CGT | 7.7 |
| ATA | 3.8 | TTA | 28.6 | GTA | 2.6 | CTA | 19.1 |
| ATC | 13.2 | TTC | 39.4 | GTC | 10.4 | CTC | 30.4 |
| ATG | 2.4 | TTG | 19.6 | GTG | 0.1 | CTG | 8.9 |
| ATT | 29.6 | TTT | 57.6 | GTT | 16.7 | CTT | 39.1 |

**SI-5: Neighbor-dependent attenuation of the damage rates**

The molecular damage rates determined above can be modified depending on their sequence context. We illustrate this effect using the example of the GPy effect, in which the formation of di-pyrimidine lesions is attenuated by a neighboring Guanine. One possible explanation is the formation of a charge-transfer state between the strong electron donor G and a pyrimidine, which suppresses damage formation[11,12]. To quantify the effect for the numerical calculations performed in SI-6, a set of tetramer damage rates are determined analogous to the trimer damage rates in SI-4.

**Table SI-t4:** Selected tetramer damage rates to approximate the effect of near-by Guanines to the formation of pyrimidine lesions.

| Tetramer | $\mu_i$ $(10^{-3}\,\frac{dmg}{base})$ | Tetramer | $\mu_i$ $(10^{-3}\,\frac{dmg}{base})$ | Tetramer | $\mu_i$ $(10^{-3}\,\frac{dmg}{base})$ |
|---|---|---|---|---|---|
| GTTG | 4 | GCCG | 2 | GCTG | 2 |
| GTTA | 13 | GCCA | 4 | GCTA | 4 |
| ATTG | 14 | ACCG | 4 | ACTG | 3 |
| ATTA | 21 | ACCA | 5 | ACTA | 5 |

Here, one considers only the tetramers that have a di-pyrimidine at the central position. For each di-pyrimidine there are edge sequences which, according to table SI-t2, do not lead to the formation of dimeric lesions within the error threshold of the method used. Examples are TA, CA or AC, AT as their order is not taken into account. As the damage rates in Table SI-t4 have already been corrected for the edge effects according to SI-2, one can directly compare, e.g. the tetramer ATTA, where the TT damage is formed without further interaction with the neighboring bases, directly with GTTA or ATTG to obtain the effect of a single Guanine as neighboring base. This reveals an attenuation of the TT damage rate by a factor of 0.7 from one neighboring Guanine ($14/21$~0.7) and 0.2 for two neighboring Guanines ($4/21{\sim}0.2$). The same result is obtained approximately for the CT or CC damage shown in table SI-t4. By this method, a matrix $AT_{W,Z}(XY)$ can now be defined that contains the attenuation factors for the damage of a dimer XY with its neighbors W and Z (WXYZ) and allows a precise calculation of UV damage for large, biased sequence pools of long DNA strands.

**Monte Carlo simulation:**

**SI-6: Numerical damage model on random DNA pools**

While the determination of UV reads in genomic sequences using high-throughput sequencing is already well established, in this work we focus on the UV damage susceptibility of sequence pools that develop starting from a strong sequence bias towards a balanced sequence space. Since there is an unmanageably large number of different such development paths, we randomly select 10000 different so called codon chronologies from these to obtain an approximate distribution of damage susceptibility per development path. Assessing the susceptibility to UV-damage of even this reduced number of developmental pathways is only possible numerically due to the large number of experiments required.

For this, we use the results from the previous chapters which yield both the molecular dimer damage rates and the effective damage rates of all other subsequences $i$ with length 3 to 8 under uniform boundary conditions. Especially for subsequences of length 4 and 6 the influence of the adjacent bases on the formation of the central dimeric lesion can now be read off to perform a numerical calculation of damage formation in large pools of random long sequences.

We implemented this simulation in a custom-made Labview program following a Monte Carlo approach (source files included), which consists of several steps:

1. Creation of the sequence pool:
   For the data shown in Figures 3 and 4, we created for each step of a codon chronology a pool of $m_{total} = 10^4$ strands, each consisting of $n = 150$ nucleotides. The strands of the sequence pool associated with the first step of a codon chronology are composed of only one specific trimer sequence and its complementary sequence. The strands of the sequence pool associated with the second step of a codon chronology are composed of the previously used trimer pair and another trimer pair. For each codon chronology, pools with an ever-larger accessible sequence space are thus created step by step until all 32 possible trimer pairs can occur in the 32nd step (see Figure 3). The individual codon chronologies differ only in the choice of specific trimer sequences that are used to randomly build up the 150mers in the respective pools. For simulation, the pool state $P$ is now stored by keeping track of all the strands including their sequence, damage state, and the number of photons absorbed for each of the bases they contain.
2. Irradiation-step of a pool in state $P$ with dose $D$ (photons/base):
   First, the photons corresponding to dose $D$ are distributed over all bases of each strand by Poisson distribution and are stored in $P$, which yields the photon number $N_{i,s,P}$ at each position $i$ of the strand $s$ in $P$.
   Since only dimeric UV lesions are considered in this work, for each dimer $XY_{s,i}$ occurring in strand $s$ at position $i$, we:
   a. Check if the dimer $XY_{s,i}$ is already damaged. If yes, we directly go to step e.
   b. determine the number of photons relevant for the formation of a lesion $NL_{s,i,P}$. Bases at the edge of the strand are considered only once, so that for $i = 1$: $NL_{1,s,P} = N_{1,s,P} + 0.5 \cdot N_{2,s,P}$, for $i = n - 1$: $NL_{n-1,s,P} = 0.5 \cdot N_{n-1,s,P} + N_{n,s,P}$ and for all other dimers $NL_{i,s,P} = 0.5 \cdot N_{i,s,P} + 0.5 \cdot N_{i+1,s,P}$.
   c. determine the contextual bases left (W) and right (Z) from the actual dimer at position $i$ and look up the context dependent damage rate using the attenuation matrix defined in SI-5 to obtain the damage probability $p_{dmg} = AT_{W,Z}(XY) \cdot \Phi_{XY}$.
   d. Execute a random generator resulting in a value $\alpha = 0..1$ and decrease $NL_{s,i,P}$ by 1. $\alpha < p_{dmg}$ confirms the damage formation, saves the change in $P$ and the algorithm directly proceeds with step e. In case of $\alpha \geq p_{dmg}$, step d is repeated until $NL_{s,i,P} = 0$.
   e. Go to the next position $i' = i + 1$ if $i < n - 1$, otherwise continue.

3. After all photons for each strand inside $P$ were evaluated and the damage states were updated in $P$, strands that contain at least one damage are considered to be "dying" strands, while strands without any damage are defined to be "surviving" strands occurring $m_{surv,P}$ times. The value "strands surviving" shown in Figure 3 and 4 accordingly is the defined as the ratio of $w_P = m_{surv,P}/m_{total}$.

The threshold to distinguish surviving and dying strands was defined as 1, because each UV lesion is assumed to be a detriment to the transfer of genetic information to the successor generation. Thus, the quantity $w_P$ reflects how many strands emerge from UV exposure without any evolutionary disadvantage. Even if the strands would suffer such a disadvantage only after several damages, this would be compensated by a slightly longer exposure i.e. higher dose. Regardless, for a constant dose, the relative susceptibility of these pools to UV irradiation can be read from the comparison of $w_P$ for different pools $P$ which exhibit different codon sequence bias.

**SI-7: Evolutionary algorithm for calculation of codon chronologies with worst and best possible UV resistance**

The procedure shown in Figure 3 is applicable for well-defined codon chronologies, i.e. when the order of all codon pairs is fixed. To test the UV sensitivity of such a chronology, 32 test pools, each containing 10000 strands of 150 bases in length, are exposed in silico to a fixed radiation dose $D$ following the procedure of the preceeding chapter. After irradiation, the number of strands without a single damage is determined for each pool, this number in relation to the number of strands initially present yields the value "strands surviving". The strands of the first of these test pools are randomly formed beforehand from the first trimer pair of the codon chronology, the strands of the second pool from the first and second trimer pairs of the codon chronology, and so on. The strands of the last test pool are formed randomly from all available codons and thus have no sequence bias. The first test pool, on the other hand, has maximum sequence bias because its strands are formed from only one trimer sequence and its complementary sequence. Each codon chronology is thus a possible evolution from a strongly biased to a completely random sequence pool.

Whereas e.g. the randomly chosen codon chronologies (gray curves in Fig 4b) are unambiguous by definition, for codon chronologies found by external criteria often no unique sequence can be specified. For example, taking the temperature stability of codon pairs as a criterion, and defining the codon chronology by sorting them by their melting temperature, some codon pairs such as GUA/UAC and CUA/UAG cannot be uniquely sorted due to their identical melting temperature.

In a chronology $O: c_1, c_2, .., (c_v, .., c_w), .., c_{32}$ of codon pairs $c_i$ this is indicated by writing ambiguously arrangeable pairs $c_v, .., c_w$ in a bracket. To be able to characterize the UV sensitivity for such a codon chronology, it makes sense to determine extremal unambiguous chronologies $O_{extr}: c_1, c_2, .., c_k, .., c_l, .. c_{32};\ c_k, .., c_l \in \{c_v, .., c_w\}$, for which the codon pairs are sorted so that they have the maximum ($O_{extr} = O_{max}$) or minimum ($O_{extr} = O_{min}$) UV sensitivity, but are still compatible with O.

For this purpose, the uniquely ordered codon pairs of a chronology are first processed as described above, i.e. for each step $t$, a sequence pool is created whose strands are randomly composed of the codon pairs $c_1,..,c_{t-1},c_t$. This is carried out until the next codon pair $c_{t+1}$ belongs to a group $(c_v,..,c_w)$ that cannot be sorted unambiguously. In this case, we will perform an $w-v$ times in-silico exposure of a pool of 150meres consisting of the codons $c_1,..,c_{t+1}$, choosing a different codon pair for $c_{t+1} \in \{c_v,..,c_w\}$ on each trial. Finally, the pool with the highest or lowest number of surviving strands is chosen, so that in each case a partially extremal codon chronology of length $t+1$: $O_{min,t+1} = c_1,..,c_{t+1,min}$ or $O_{max,t+1} = c_1,..,c_{t+1,max}$ can be determined. For the next step $t+2$, the same procedure is repeated for the codon group $(c_v,..,c_w)$ reduced by $c_{t+1}$. This evolutionary algorithm finally yields the two codon chronologies

$O_{min} = c_1,..,c_{t+1,min},..,c_{t+w-v,min},..c_w$ or $O_{max} = c_1,..,c_{t+1,max},..,c_{t+w-v,max},..c_w$, which represent the chronologies compatible with $O$ with maximum or minimum UV sensitivity. If several groups of non-arrangeable codon pairs are present, they are processed in an analogous manner one after the other. $O_{min}$ and $O_{max}$ are thus the limit cases of the UV sensitivity of a codon chronology whose codon pairs are partially not uniquely arrangeable according to external criteria.

A special case will be treated here in particular, namely when there is no external criterion and thus all codon pairs can occur in any order. In this case, the above procedure results in the sequences

$O_{min} = c_{1,min},..,c_{32,min}$ and $O_{max} = c_{1,max},..,c_{32,max}$ , which represent the optimal or worst development of the sequence bias in prebiotic pools purely under UV radiation as the only selection pressure. The sequence $O_{min}$ is explicitly shown in Table 1, the sequence $O_{max}$ is shown in Table SI-t5.

**SI-t5** Worst case codon chronology with UV as only selection pressure. Strands surviving is shown as red line in Figure 4b.

| Rank | Codon pair | | Rank | Codon pair | |
|---|---|---|---|---|---|
| 1 | TTT | AAA | 17 | TCG | CGA |
| 2 | GAA | TTC | 18 | GGG | CCC |
| 3 | CTT | AAG | 19 | CCA | TGG |
| 4 | TTA | TAA | 20 | ATA | TAT |
| 5 | ATT | AAT | 21 | GAC | GTC |
| 6 | TCT | AGA | 22 | ACA | TGT |
| 7 | CAA | TTG | 23 | GGT | ACC |
| 8 | GTT | AAC | 24 | CTG | CAG |
| 9 | GGA | TCC | 25 | ATG | CAT |
| 10 | CCT | AGG | 26 | GTA | TAC |
| 11 | ACT | AGT | 27 | GTG | CAC |
| 12 | TCA | TGA | 28 | CCG | CGG |
| 13 | GAG | CTC | 29 | GGC | GCC |
| 14 | GAT | ATC | 30 | ACG | CGT |
| 15 | CTA | TAG | 31 | GCG | CGC |
| 16 | GCT | AGC | 32 | GCA | TGC |

To determine the statistical error in Table 1, the in silico exposures are repeated three times using the above procedure to find the mean and standard deviation for the strands surviving at each step. If the standard deviation is greater than the difference in UV sensitivity of the test pools with two different codon pairs when determining the codon pair for the next step, these codon pairs are given the same rank. This means that for codon pairs with the same rank, with respect to UV exposure, it does not matter in which order they are in the codon chronology.

**SI-8: Deduction of codon chronologies from amino acid chronologies**

Often in the literature, not codons but amino acid chronologies are given, i.e. the sequence of amino acids according to their predicted occurrence in evolution[13]. While it is not clear whether the canonical genetic code was already fixed in this early phase of evolution, these amino acid chronologies provide a possible starting point for the sequence bias of the "proto-genomes", i.e. the genomes of the life forms existing at that time. It cannot be ruled out that other codon sequences in the protogenome were used for the restricted set of amino acids at an early stage. However, it is plausible that before lifeforms started to use additional amino acids, their then newly associated codon sequences had already been used before in a different way compared to the codon sequences associated to the already established amino acids. The derivation of codon chronologies from amino acid sequences is thus an approximation for the evolution of sequence bias for protogenomes.

For the construction of codon chronologies $O: c_1, \ldots, c_{32}$ from amino acid chronologies $OA: a_1, \ldots, a_{21}$, an analogous procedure was used to determine the consensus order for codons[13]. For each amino acid $a$ of $OA$, we:

1. determine all codons for the amino acid $a$ and their reverse complementary sequences according to the canonical genetic code.
2. choose from these codon pairs the pair that can be obtained from the already used codon pairs with the change of only one base.
3. If there are several corresponding codon pairs with this property, we choose the codon pair with the highest thermal stability from this group.

This is done until all 32 codon pairs have been arranged. A possible late rearrangement, e.g. by codon capture, is not taken into account. This is plausible as the selective effect of UV radiation is expected to play a role especially for less evolved life forms that had only a limited set of amino acids available.

The codon sequences used in Figure 4c were assigned a "criteria index" and explicitly shown in Table SI-t6 with their corresponding amino acid sequence. Should amino acids $a_v, .., a_w$ of a chronology not be unambiguously sortable, they are grouped in brackets: $AO: a_1, \ldots, (a_v, .., a_w), .. a_{21}$. This is taken into account in the determination of the codon sequences through the commutability of the associated codon pairs and results in two limit codon chronologies with maximal and minimal UV sensitivity.

**SI-t6** Amino acid chronologies[13] and their calculated worst and best case codon chronologies.

| Criterion index | 1 | 2 | 3 | 4 | |
|---|---|---|---|---|---|
| Reference | Trifonov[13] | Haig[14], Wong[15], Papentin[16], Dufton[17] | Arques[18], Brooks[19,20], Eck[21] | Xia[22] | |
| AA chronology | GADVPSETLR NIQHKCFYMW | GASPVCDTNLKIE | AGVSDLTPEIR KNQHFMYCW | AGSPRDTCE (VW)HL(MQ)INYFK | |
| UV sensitivity | unambiguous | unambiguous | unambiguous | min | max |
| Codon chronology | GGC,GCC | GGC,GCC | GGC,GCC | GGC,GCC | GGC,GCC |
| | GAC,GTC | GGA,TCC | GAC,GTC | GGA,TCC | GGA,TCC |
| | GGG,CCC | GGG,CCC | GGA,TCC | GGG,CCC | GGG,CCC |
| | GGA,TCC | GCT,AGC | GAG,CTC | GCG,CGC | GCG,CGC |
| | GCT,AGC | GAC,GTC | GGT,ACC | GCT,AGC | GCT,AGC |
| | GAG,CTC | GCA,TGC | GGG,CCC | GAC,GTC | GAC,GTC |
| | GGT,ACC | GGT,ACC | TCA,TGA | GGT,ACC | GGT,ACC |
| | GCG,CGC | TCA,TGA | ACT,AGT | GCA,TGC | GCA,TGC |
| | CCG,CGG | GTT,AAC | ACA,TGT | CCG,CGG | CCG,CGG |
| | CCT,AGG | GAG,CTC | GAA,TTC | CCT,AGG | CCT,AGG |
| | TCG,CGA | CTT,AAG | GAT,ATC | GAG,CTC | GAG,CTC |
| | TCA,TGA | ACT,AGT | GCG,CGC | GTG,CAC | CCA,TGG |
| | TCT,AGA | GAT,ATC | CTT,AAG | CCA,TGG | GTG,CAC |
| | ACG,CGT | ACA,TGT | GCT,AGC | ATG,CAT | ATG,CAT |
| | ACT,AGT | GAA,TTC | GTG,CAC | TCG,CGA | TCG,CGA |
| | ACA,TGT | ATG,CAT | CCG,CGG | GTT,AAC | GTT,AAC |
| | GTT,AAC | GCG,CGC | TTT,AAA | TCA,TGA | TCA,TGA |
| | GAT,ATC | TTT,AAA | CCT,AGG | TCT,AGA | TCT,AGA |
| | GTG,CAC | CCG,CGG | ATG,CAT | ACG,CGT | ACG,CGT |
| | CTT,AAG | GTA,TAC | GTA,TAC | ACT,AGT | ACT,AGT |
| | GCA,TGC | CCT,AGG | GCA,TGC | ACA,TGT | ACA,TGT |
| | GAA,TTC | CCA,TGG | CCA,TGG | GAT,ATC | GAT,ATC |
| | CTG,CAG | TCG,CGA | TCG,CGA | CTG,CAG | CTG,CAG |
| | GTA,TAC | TCT,AGA | TCT,AGA | GTA,TAC | GTA,TAC |
| | ATG,CAT | ACG,CGT | ACG,CGT | GAA,TTC | GAA,TTC |
| | CCA,TGG | GTG,CAC | CTG,CAG | CTA,TAG | CTA,TAG |
| | CTA,TAG | CTG,CAG | CTA,TAG | CAA,TTG | CAA,TTG |
| | CAA,TTG | CTA,TAG | GTT,AAC | CTT,AAG | CTT,AAG |
| | ATA,TAT | CAA,TTG | CAA,TTG | ATA,TAT | ATA,TAT |
| | ATT,AAT | ATA,TAT | ATA,TAT | ATT,AAT | ATT,AAT |
| | TTA,TAA | ATT,AAT | ATT,AAT | TTA,TAA | TTA,TAA |
| | TTT,AAA | TTA,TAA | TTA,TAA | TTT,AAA | TTT,AAA |

| 5 | | 6 | | 7 | | 8 | |
|---|---|---|---|---|---|---|---|
| Osawa[23] | | Crick[24] | | Eigen[25,26] | | Ferreira[27] | |
| (ADEFGHP)VSTLRI Q(KNY)(MW)C | | (DGNS)(ACEFHIK LMPQRTVWY) | | (AG)(DINSTV)(CE FHKLMPQRWY) | | (FGKLNP)(CDEH QRSTVW)(AIMY) | |
| min | max | min | max | min | max | min | max |
| GTG,CAC | GAA,TTC | GGC,GCC | GGA,TCC | GGC,GCC | GGC,GCC | GGC,GCC | TTT,AAA |
| GGC,GCC | GAG,CTC | GAC,GTC | GTT,AAC | GGT,ACC | GGA,TCC | GGG,CCC | GAA,TTC |
| GAC,GTC | GGG,CCC | GTT,AAC | GAC,GTC | GAC,GTC | GAT,ATC | GAA,TTC | GGG,CCC |
| GGG,CCC | GAC,GTC | GGA,TCC | GGC,GCC | GTT,AAC | GTT,AAC | TTT,AAA | GGC,GCC |
| GAG,CTC | GGC,GCC | GCG,CGC | GAA,TTC | GAT,ATC | GAC,GTC | GCG,CGC | GGA,TCC |
| GAA,TTC | GTG,CAC | GTG,CAC | CTT,AAG | GGA,TCC | GGT,ACC | GTG,CAC | GAC,GTC |
| GGA,TCC | GGA,TCC | GCA,TGC | GAG,CTC | GCG,CGC | GAA,TTC | GCA,TGC | CCA,TGG |
| GCT,AGC | GCT,AGC | ATG,CAT | GAT,ATC | GTG,CAC | CTT,AAG | CCA,TGG | GGT,ACC |
| GGT,ACC | GGT,ACC | GTA,TAC | CCA,TGG | ATG,CAT | GAG,CTC | GGT,ACC | GCA,TGC |
| CTG,CAG | CTG,CAG | GGT,ACC | GGG,CCC | GCA,TGC | GGG,CCC | GAC,GTC | GTG,CAC |
| GCG,CGC | GCG,CGC | CCA,TGG | GGT,ACC | GTA,TAC | CCA,TGG | GGA,TCC | GCG,CGC |
| GAT,ATC | GAT,ATC | GGG,CCC | GTA,TAC | CCA,TGG | GTA,TAC | GCT,AGC | GCT,AGC |
| CCG,CGG | CCG,CGG | GAT,ATC | GCA,TGC | GGG,CCC | GTG,CAC | CCG,CGG | CCG,CGG |
| ATG,CAT | ATG,CAT | GAG,CTC | GTG,CAC | GAG,CTC | GCA,TGC | CCT,AGG | CCT,AGG |
| CCT,AGG | CCT,AGG | GAA,TTC | ATG,CAT | CTT,AAG | GCG,CGC | TCG,CGA | TCG,CGA |
| GTA,TAC | CTT,AAG | CTT,AAG | GCG,CGC | GAA,TTC | ATG,CAT | GAG,CTC | GAG,CTC |
| CTT,AAG | GTA,TAC | GCT,AGC | GCT,AGC | GCT,AGC | GCT,AGC | TCA,TGA | TCA,TGA |
| CCA,TGG | CCA,TGG | CCG,CGG | CCG,CGG | CCG,CGG | CCG,CGG | TCT,AGA | TCT,AGA |
| GCA,TGC | GCA,TGC | CCT,AGG | CCT,AGG | CCT,AGG | CCT,AGG | ACG,CGT | ACG,CGT |
| TCG,CGA | TCG,CGA | TCG,CGA | TCG,CGA | TCG,CGA | TCG,CGA | ACT,AGT | ACT,AGT |
| TCA,TGA | TCA,TGA | TCA,TGA | TCA,TGA | TCA,TGA | TCA,TGA | ACA,TGT | ACA,TGT |
| TCT,AGA | TCT,AGA | TCT,AGA | TCT,AGA | TCT,AGA | TCT,AGA | ATG,CAT | GAT,ATC |
| ACG,CGT | ACG,CGT | ACG,CGT | ACG,CGT | ACG,CGT | ACG,CGT | GTA,TAC | GTA,TAC |
| ACT,AGT | ACT,AGT | ACT,AGT | ACT,AGT | ACT,AGT | ACT,AGT | GAT,ATC | ATG,CAT |
| ACA,TGT | ACA,TGT | ACA,TGT | ACA,TGT | ACA,TGT | ACA,TGT | CTG,CAG | CTG,CAG |
| CTA,TAG | CTA,TAG | CTG,CAG | CTG,CAG | CTG,CAG | CTG,CAG | CTA,TAG | CTA,TAG |
| GTT,AAC | GTT,AAC | CTA,TAG | CTA,TAG | CTA,TAG | CTA,TAG | GTT,AAC | GTT,AAC |
| CAA,TTG | CAA,TTG | CAA,TTG | CAA,TTG | CAA,TTG | CAA,TTG | CAA,TTG | CAA,TTG |
| ATA,TAT | ATA,TAT | ATA,TAT | ATA,TAT | ATA,TAT | ATA,TAT | CTT,AAG | CTT,AAG |
| ATT,AAT | ATT,AAT | ATT,AAT | ATT,AAT | ATT,AAT | ATT,AAT | ATA,TAT | ATA,TAT |
| TTA,TAA | TTA,TAA | TTA,TAA | TTA,TAA | TTA,TAA | TTA,TAA | ATT,AAT | ATT,AAT |
| TTT,AAA | TTT,AAA | TTT,AAA | TTT,AAA | TTT,AAA | TTT,AAA | TTA,TAA | TTA,TAA |

| 9 | | 10 | | 11 | | 12 | |
|---|---|---|---|---|---|---|---|
| Nelsestuen[28] | | Möller[29] | | Chaley[30] | | Henikoff[31] | |
| (DEFHIKLMSTVY) (ACGNPQRW) | | (ADGV)(CEFHIK LMNPQRSTWY) | | QHP(LS)GCWR V(DE)AYT(IM)F(KN) | | (AILSV)(EKMQRT) (DFGN)(PY)HCW | |
| min | max | min | max | min | max | min | max |
| GTG,CAC | CTT,AAG | GAC,GTC | GGC,GCC | GTG,CAC | GTG,CAC | GAC,GTC | GAG,CTC |
| ATG,CAT | GAA,TTC | GGC,GCC | GAC,GTC | GGG,CCC | GGG,CCC | GGC,GCC | GGA,TCC |
| GTA,TAC | GAG,CTC | GCG,CGC | GAA,TTC | CTG,CAG | GGA,TCC | GAT,ATC | GAT,ATC |
| GGT,ACC | GGA,TCC | GTG,CAC | CTT,AAG | GGA,TCC | CTG,CAG | GGA,TCC | GAC,GTC |
| GAC,GTC | GAT,ATC | ATG,CAT | GAG,CTC | GCA,TGC | GCA,TGC | GAG,CTC | GGC,GCC |
| GAT,ATC | GAC,GTC | GCA,TGC | GGA,TCC | CCA,TGG | CCA,TGG | GCG,CGC | CTT,AAG |
| GGA,TCC | GGT,ACC | GTA,TAC | GAT,ATC | GCG,CGC | GCG,CGC | ATG,CAT | GGT,ACC |
| GAG,CTC | GTA,TAC | GGT,ACC | GGG,CCC | GAC,GTC | GAC,GTC | GGT,ACC | ATG,CAT |
| GAA,TTC | GTG,CAC | CCA,TGG | GGT,ACC | GAG,CTC | GAG,CTC | CTT,AAG | GCG,CGC |
| CTT,AAG | ATG,CAT | GGG,CCC | CCA,TGG | GGC,GCC | GGC,GCC | GAA,TTC | GAA,TTC |
| GCG,CGC | GGG,CCC | GAT,ATC | GTA,TAC | GTA,TAC | GTA,TAC | GTA,TAC | GGG,CCC |
| GCA,TGC | CCA,TGG | GGA,TCC | ATG,CAT | GCT,AGC | GCT,AGC | GGG,CCC | GTA,TAC |
| GGC,GCC | GGC,GCC | GAG,CTC | GCA,TGC | GGT,ACC | GGT,ACC | GTG,CAC | GTG,CAC |
| CCA,TGG | GCG,CGC | GAA,TTC | GCG,CGC | CCG,CGG | CCG,CGG | GCT,AGC | GCT,AGC |
| GGG,CCC | GCA,TGC | CTT,AAG | GTG,CAC | CCT,AGG | CCT,AGG | GCA,TGC | GCA,TGC |
| GCT,AGC | GCT,AGC | GCT,AGC | GCT,AGC | TCG,CGA | TCG,CGA | CCG,CGG | CCG,CGG |
| CCG,CGG | CCG,CGG | CCG,CGG | CCG,CGG | TCA,TGA | TCA,TGA | CCT,AGG | CCT,AGG |
| CCT,AGG | CCT,AGG | CCT,AGG | CCT,AGG | TCT,AGA | TCT,AGA | CCA,TGG | CCA,TGG |
| TCG,CGA | TCG,CGA | TCG,CGA | TCG,CGA | ACG,CGT | ACG,CGT | TCG,CGA | TCG,CGA |
| TCA,TGA | TCA,TGA | TCA,TGA | TCA,TGA | ACT,AGT | ACT,AGT | TCA,TGA | TCA,TGA |
| TCT,AGA | TCT,AGA | TCT,AGA | TCT,AGA | ACA,TGT | ACA,TGT | TCT,AGA | TCT,AGA |
| ACG,CGT | ACG,CGT | ACG,CGT | ACG,CGT | ATG,CAT | GAT,ATC | ACG,CGT | ACG,CGT |
| ACT,AGT | ACT,AGT | ACT,AGT | ACT,AGT | GAT,ATC | ATG,CAT | ACT,AGT | ACT,AGT |
| ACA,TGT | ACA,TGT | ACA,TGT | ACA,TGT | GAA,TTC | GAA,TTC | ACA,TGT | ACA,TGT |
| CTG,CAG | CTG,CAG | CTG,CAG | CTG,CAG | CTA,TAG | CTA,TAG | CTG,CAG | CTG,CAG |
| CTA,TAG | CTA,TAG | CTA,TAG | CTA,TAG | CTT,AAG | CTT,AAG | CTA,TAG | CTA,TAG |
| GTT,AAC | GTT,AAC | GTT,AAC | GTT,AAC | GTT,AAC | GTT,AAC | GTT,AAC | GTT,AAC |
| CAA,TTG | CAA,TTG | CAA,TTG | CAA,TTG | CAA,TTG | CAA,TTG | CAA,TTG | CAA,TTG |
| ATA,TAT | ATA,TAT | ATA,TAT | ATA,TAT | ATA,TAT | ATA,TAT | ATA,TAT | ATA,TAT |
| ATT,AAT | ATT,AAT | ATT,AAT | ATT,AAT | ATT,AAT | ATT,AAT | ATT,AAT | ATT,AAT |
| TTA,TAA | TTA,TAA | TTA,TAA | TTA,TAA | TTA,TAA | TTA,TAA | TTA,TAA | TTA,TAA |
| TTT,AAA | TTT,AAA | TTT,AAA | TTT,AAA | TTT,AAA | TTT,AAA | TTT,AAA | TTT,AAA |

| 13 | | 14 | | 15 | | 16 | |
|---|---|---|---|---|---|---|---|
| Eigen[32] | | Kvenvolden[33] | | Riddle[34] | | Arques[18] | |
| V(AGP)(ENRT)(LQS) (CDFHIKMYW) | | (AG)(DEPV)(CFHIK LMNQRSTWY) | | (AGEIK)(CDFHLM NPQRSTVWY) | | (ADEFGIKLNQTVY) (CHMPRSW) | |
| min | max | min | max | min | max | min | max |
| GAC,GTC | GAC,GTC | GGC,GCC | GGC,GCC | GGC,GCC | CTT,AAG | GTG,CAC | CTT,AAG |
| GGC,GCC | GGG,CCC | GAC,GTC | GAG,CTC | GAT,ATC | GAG,CTC | GTA,TAC | GAA,TTC |
| GGG,CCC | GGC,GCC | GGG,CCC | GGG,CCC | GAG,CTC | GAT,ATC | GGC,GCC | GAG,CTC |
| GCG,CGC | GAG,CTC | GAG,CTC | GAC,GTC | CTT,AAG | GGC,GCC | GGT,ACC | GAT,ATC |
| GGT,ACC | GTT,AAC | GTG,CAC | GAA,TTC | GCG,CGC | GAA,TTC | GAC,GTC | GGT,ACC |
| GTT,AAC | GGT,ACC | GCG,CGC | CTT,AAG | ATG,CAT | GGA,TCC | GAT,ATC | GAC,GTC |
| GAG,CTC | GCG,CGC | ATG,CAT | GGA,TCC | GCA,TGC | GGG,CCC | GAG,CTC | GTA,TAC |
| GTG,CAC | GGA,TCC | GCA,TGC | GAT,ATC | GTA,TAC | GAC,GTC | CTT,AAG | GGC,GCC |
| GGA,TCC | GTG,CAC | GTA,TAC | CCA,TGG | GGT,ACC | GGT,ACC | GAA,TTC | GTG,CAC |
| GCT,AGC | GCT,AGC | CCA,TGG | GGT,ACC | GAC,GTC | CCA,TGG | GCG,CGC | GGA,TCC |
| GCA,TGC | CTT,AAG | GGT,ACC | GTA,TAC | CCA,TGG | GTA,TAC | GCA,TGC | GGG,CCC |
| ATG,CAT | GAA,TTC | GAT,ATC | GCA,TGC | GGG,CCC | GCG,CGC | ATG,CAT | CCA,TGG |
| GTA,TAC | GAT,ATC | GGA,TCC | ATG,CAT | GGA,TCC | ATG,CAT | CCA,TGG | GCA,TGC |
| CCA,TGG | CCA,TGG | GAA,TTC | GTG,CAC | GAA,TTC | GCA,TGC | GGG,CCC | ATG,CAT |
| GAT,ATC | GCA,TGC | CTT,AAG | GCG,CGC | GCT,AGC | GCT,AGC | GGA,TCC | GCG,CGC |
| GAA,TTC | GTA,TAC | GCT,AGC | GCT,AGC | CCG,CGG | CCG,CGG | GCT,AGC | GCT,AGC |
| CTT,AAG | ATG,CAT | CCG,CGG | CCG,CGG | CCT,AGG | CCT,AGG | CCG,CGG | CCG,CGG |
| CCG,CGG | CCG,CGG | CCT,AGG | CCT,AGG | TCG,CGA | TCG,CGA | CCT,AGG | CCT,AGG |
| CCT,AGG | CCT,AGG | TCG,CGA | TCG,CGA | TCA,TGA | TCA,TGA | TCG,CGA | TCG,CGA |
| TCG,CGA | TCG,CGA | TCA,TGA | TCA,TGA | TCT,AGA | TCT,AGA | TCA,TGA | TCA,TGA |
| TCA,TGA | TCA,TGA | TCT,AGA | TCT,AGA | ACG,CGT | ACG,CGT | TCT,AGA | TCT,AGA |
| TCT,AGA | TCT,AGA | ACG,CGT | ACG,CGT | ACT,AGT | ACT,AGT | ACG,CGT | ACG,CGT |
| ACG,CGT | ACG,CGT | ACT,AGT | ACT,AGT | ACA,TGT | ACA,TGT | ACT,AGT | ACT,AGT |
| ACT,AGT | ACT,AGT | ACA,TGT | ACA,TGT | GTG,CAC | GTG,CAC | ACA,TGT | ACA,TGT |
| ACA,TGT | ACA,TGT | CTG,CAG | CTG,CAG | CTG,CAG | CTG,CAG | CTG,CAG | CTG,CAG |
| CTG,CAG | CTG,CAG | CTA,TAG | CTA,TAG | CTA,TAG | CTA,TAG | CTA,TAG | CTA,TAG |
| CTA,TAG | CTA,TAG | GTT,AAC | GTT,AAC | GTT,AAC | GTT,AAC | GTT,AAC | GTT,AAC |
| CAA,TTG | CAA,TTG | CAA,TTG | CAA,TTG | CAA,TTG | CAA,TTG | CAA,TTG | CAA,TTG |
| ATA,TAT | ATA,TAT | ATA,TAT | ATA,TAT | ATA,TAT | ATA,TAT | ATA,TAT | ATA,TAT |
| ATT,AAT | ATT,AAT | ATT,AAT | ATT,AAT | ATT,AAT | ATT,AAT | ATT,AAT | ATT,AAT |
| TTA,TAA | TTA,TAA | TTA,TAA | TTA,TAA | TTA,TAA | TTA,TAA | TTA,TAA | TTA,TAA |
| TTT,AAA | TTT,AAA | TTT,AAA | TTT,AAA | TTT,AAA | TTT,AAA | TTT,AAA | TTT,AAA |

| 17 | | 18 | | 19 | | 20 | |
|---|---|---|---|---|---|---|---|
| Yarus[35] | | Jukes[36] | | Wong[37] | | Harman[38] | |
| R(ACDEFGHIKL MNPQSTVWY) | | (ADEGHLPQRV) (CFIKNSTY)(MW) | | (ADEGS)V(PT)(IL) FCY(KR)(NQ)H(MW) | | GPAR(DENQST)(H K)C(FILVY)(MW) | |
| min | max | min | max | min | max | min | max |
| GCG,CGC | GCG,CGC | GTG,CAC | GAG,CTC | GGC,GCC | GGA,TCC | GGC,GCC | GGC,GCC |
| GCA,TGC | CTT,AAG | GCG,CGC | GAC,GTC | GAC,GTC | GAG,CTC | GGG,CCC | GGG,CCC |
| GTG,CAC | GAA,TTC | GGC,GCC | GGG,CCC | GGA,TCC | GAC,GTC | GCG,CGC | GCG,CGC |
| ATG,CAT | GAG,CTC | GAC,GTC | GGC,GCC | GAG,CTC | GGC,GCC | GTG,CAC | GAG,CTC |
| GTA,TAC | GGA,TCC | GGG,CCC | GCG,CGC | GGT,ACC | GGG,CCC | GGT,ACC | GGA,TCC |
| GGC,GCC | GAT,ATC | GAG,CTC | GTG,CAC | GGG,CCC | GGT,ACC | GAC,GTC | GTT,AAC |
| CCA,TGG | GGG,CCC | GCA,TGC | GAA,TTC | GCT,AGC | GCT,AGC | GTT,AAC | GAC,GTC |
| GGT,ACC | GAC,GTC | GTA,TAC | CTT,AAG | TCA,TGA | TCA,TGA | GGA,TCC | GGT,ACC |
| GAC,GTC | GGT,ACC | GGT,ACC | GGA,TCC | ACT,AGT | ACT,AGT | GAG,CTC | GTG,CAC |
| GGG,CCC | CCA,TGG | GAT,ATC | GAT,ATC | ACA,TGT | ACA,TGT | GCT,AGC | GCT,AGC |
| GAT,ATC | GTA,TAC | GGA,TCC | GTA,TAC | CTG,CAG | GAT,ATC | CTT,AAG | CTT,AAG |
| GGA,TCC | GCA,TGC | CTT,AAG | GGT,ACC | GAT,ATC | CTG,CAG | GCA,TGC | GCA,TGC |
| GAG,CTC | GGC,GCC | GAA,TTC | GCA,TGC | GAA,TTC | GAA,TTC | GTA,TAC | GAA,TTC |
| GAA,TTC | ATG,CAT | GCT,AGC | GCT,AGC | GCA,TGC | GCA,TGC | GAT,ATC | GAT,ATC |
| CTT,AAG | GTG,CAC | CCG,CGG | CCG,CGG | GTA,TAC | GTA,TAC | GAA,TTC | GTA,TAC |
| GCT,AGC | GCT,AGC | CCT,AGG | CCT,AGG | GCG,CGC | CTT,AAG | CCG,CGG | CCG,CGG |
| CCG,CGG | CCG,CGG | ATG,CAT | CCA,TGG | CTT,AAG | GCG,CGC | ATG,CAT | CCA,TGG |
| CCT,AGG | CCT,AGG | CCA,TGG | ATG,CAT | GTG,CAC | GTG,CAC | CCA,TGG | ATG,CAT |
| TCG,CGA | TCG,CGA | TCG,CGA | TCG,CGA | CCG,CGG | CCG,CGG | CCT,AGG | CCT,AGG |
| TCA,TGA | TCA,TGA | TCA,TGA | TCA,TGA | ATG,CAT | CCA,TGG | TCG,CGA | TCG,CGA |
| TCT,AGA | TCT,AGA | TCT,AGA | TCT,AGA | CCA,TGG | ATG,CAT | TCA,TGA | TCA,TGA |
| ACG,CGT | ACG,CGT | ACG,CGT | ACG,CGT | CCT,AGG | CCT,AGG | TCT,AGA | TCT,AGA |
| ACT,AGT | ACT,AGT | ACT,AGT | ACT,AGT | TCG,CGA | TCG,CGA | ACG,CGT | ACG,CGT |
| ACA,TGT | ACA,TGT | ACA,TGT | ACA,TGT | TCT,AGA | TCT,AGA | ACT,AGT | ACT,AGT |
| CTG,CAG | CTG,CAG | CTG,CAG | CTG,CAG | ACG,CGT | ACG,CGT | ACA,TGT | ACA,TGT |
| CTA,TAG | CTA,TAG | CTA,TAG | CTA,TAG | CTA,TAG | CTA,TAG | CTG,CAG | CTG,CAG |
| GTT,AAC | GTT,AAC | GTT,AAC | GTT,AAC | GTT,AAC | GTT,AAC | CTA,TAG | CTA,TAG |
| CAA,TTG | CAA,TTG | CAA,TTG | CAA,TTG | CAA,TTG | CAA,TTG | CAA,TTG | CAA,TTG |
| ATA,TAT | ATA,TAT | ATA,TAT | ATA,TAT | ATA,TAT | ATA,TAT | ATA,TAT | ATA,TAT |
| ATT,AAT | ATT,AAT | ATT,AAT | ATT,AAT | ATT,AAT | ATT,AAT | ATT,AAT | ATT,AAT |
| TTA,TAA | TTA,TAA | TTA,TAA | TTA,TAA | TTA,TAA | TTA,TAA | TTA,TAA | TTA,TAA |
| TTT,AAA | TTT,AAA | TTT,AAA | TTT,AAA | TTT,AAA | TTT,AAA | TTT,AAA | TTT,AAA |

| 21 | | 22 | | 23 | | 24 | |
|---|---|---|---|---|---|---|---|
| N16 in Trifonov[13] | | Yarus[35] | | Taylor[39] | | Ikehara[40] | |
| (ADEGKRSTV) (CFHILMNPQWY) | | (ADEGV)(CFHIK LMNPQRSTWY) | | (ADGV)(LPR)(CIK QST)(EFHMNWY) | | (ADGV)E(HLPQR) (CFIKMNSTWY) | |
| min | max | min | max | min | max | min | max |
| GCG,CGC | CTT,AAG | GGC,GCC | GAG,CTC | GAC,GTC | GGC,GCC | GAC,GTC | GGC,GCC |
| GGC,GCC | GAG,CTC | GAC,GTC | GAC,GTC | GGC,GCC | GAC,GTC | GGC,GCC | GAC,GTC |
| GGT,ACC | GGA,TCC | GAG,CTC | GGC,GCC | GCG,CGC | GAG,CTC | GAG,CTC | GAG,CTC |
| GAC,GTC | GGT,ACC | GCG,CGC | GAA,TTC | GGG,CCC | GGG,CCC | GCG,CGC | GGG,CCC |
| GGA,TCC | GAC,GTC | GTG,CAC | CTT,AAG | GAG,CTC | GCG,CGC | GTG,CAC | GCG,CGC |
| GAG,CTC | GGC,GCC | GCA,TGC | GGA,TCC | GTG,CAC | CTT,AAG | GGG,CCC | GTG,CAC |
| CTT,AAG | GCG,CGC | ATG,CAT | GAT,ATC | GCA,TGC | GGA,TCC | GCA,TGC | CTT,AAG |
| GTG,CAC | GAA,TTC | GTA,TAC | GGG,CCC | GGT,ACC | GAT,ATC | ATG,CAT | GAA,TTC |
| ATG,CAT | GAT,ATC | CCA,TGG | CCA,TGG | GAT,ATC | GGT,ACC | GTA,TAC | GGA,TCC |
| GCA,TGC | GGG,CCC | GGT,ACC | GGT,ACC | GGA,TCC | GCA,TGC | CCA,TGG | GAT,ATC |
| GTA,TAC | GTA,TAC | GGG,CCC | GTA,TAC | CTT,AAG | GTG,CAC | GGT,ACC | CCA,TGG |
| CCA,TGG | CCA,TGG | GAT,ATC | ATG,CAT | GCT,AGC | GCT,AGC | GAT,ATC | GGT,ACC |
| GGG,CCC | GCA,TGC | GGA,TCC | GCA,TGC | CCG,CGG | CCG,CGG | GGA,TCC | GTA,TAC |
| GAT,ATC | GTG,CAC | GAA,TTC | GTG,CAC | CCT,AGG | CCT,AGG | GAA,TTC | GCA,TGC |
| GAA,TTC | ATG,CAT | CTT,AAG | GCG,CGC | ATG,CAT | GAA,TTC | CTT,AAG | ATG,CAT |
| GCT,AGC | GCT,AGC | GCT,AGC | GCT,AGC | GTA,TAC | CCA,TGG | GCT,AGC | GCT,AGC |
| CCG,CGG | CCG,CGG | CCG,CGG | CCG,CGG | CCA,TGG | GTA,TAC | CCG,CGG | CCG,CGG |
| CCT,AGG | CCT,AGG | CCT,AGG | CCT,AGG | GAA,TTC | ATG,CAT | CCT,AGG | CCT,AGG |
| TCG,CGA | TCG,CGA | TCG,CGA | TCG,CGA | TCG,CGA | TCG,CGA | TCG,CGA | TCG,CGA |
| TCA,TGA | TCA,TGA | TCA,TGA | TCA,TGA | TCA,TGA | TCA,TGA | TCA,TGA | TCA,TGA |
| TCT,AGA | TCT,AGA | TCT,AGA | TCT,AGA | TCT,AGA | TCT,AGA | TCT,AGA | TCT,AGA |
| ACG,CGT | ACG,CGT | ACG,CGT | ACG,CGT | ACG,CGT | ACG,CGT | ACG,CGT | ACG,CGT |
| ACT,AGT | ACT,AGT | ACT,AGT | ACT,AGT | ACT,AGT | ACT,AGT | ACT,AGT | ACT,AGT |
| ACA,TGT | ACA,TGT | ACA,TGT | ACA,TGT | ACA,TGT | ACA,TGT | ACA,TGT | ACA,TGT |
| CTG,CAG | CTG,CAG | CTG,CAG | CTG,CAG | CTG,CAG | CTG,CAG | CTG,CAG | CTG,CAG |
| CTA,TAG | CTA,TAG | CTA,TAG | CTA,TAG | CTA,TAG | CTA,TAG | CTA,TAG | CTA,TAG |
| GTT,AAC | GTT,AAC | GTT,AAC | GTT,AAC | GTT,AAC | GTT,AAC | GTT,AAC | GTT,AAC |
| CAA,TTG | CAA,TTG | CAA,TTG | CAA,TTG | CAA,TTG | CAA,TTG | CAA,TTG | CAA,TTG |
| ATA,TAT | ATA,TAT | ATA,TAT | ATA,TAT | ATA,TAT | ATA,TAT | ATA,TAT | ATA,TAT |
| ATT,AAT | ATT,AAT | ATT,AAT | ATT,AAT | ATT,AAT | ATT,AAT | ATT,AAT | ATT,AAT |
| TTA,TAA | TTA,TAA | TTA,TAA | TTA,TAA | TTA,TAA | TTA,TAA | TTA,TAA | TTA,TAA |
| TTT,AAA | TTT,AAA | TTT,AAA | TTT,AAA | TTT,AAA | TTT,AAA | TTT,AAA | TTT,AAA |

| 25 | | 26 | | 27 | | 28 | |
|---|---|---|---|---|---|---|---|
| N54 in Trifonov[13] | | N55 in Trifonov[13] | | N57 in Trifonov[13] | | Baumann[41] | |
| (GPS)(DEFKLN) (AHQRV)(CIMTWY) | | GSDNK(AF)(HT)E QL(PV)R(CW)(IM)Y | | G(DS)A(LPV)(EFH IKMNQR)(CTWY) | | (ADEGILPQRSTV) (KN)(CFHY)(MW) | |
| min | max | min | max | min | max | min | max |
| GGC,GCC | GGA,TCC | GGC,GCC | GGC,GCC | GGC,GCC | GGC,GCC | GTG,CAC | GAG,CTC |
| GGG,CCC | GGG,CCC | GGA,TCC | GGA,TCC | GAC,GTC | GGA,TCC | GCG,CGC | GGA,TCC |
| GGA,TCC | GGC,GCC | GCT,AGC | GCT,AGC | GGA,TCC | GAC,GTC | GGC,GCC | GAT,ATC |
| GAC,GTC | CTT,AAG | GAC,GTC | GAC,GTC | GGG,CCC | GAG,CTC | GGT,ACC | GGG,CCC |
| GAG,CTC | GAA,TTC | TCA,TGA | TCA,TGA | GAG,CTC | GGG,CCC | GAC,GTC | GGT,ACC |
| CTT,AAG | GAG,CTC | GTT,AAC | GTT,AAC | GTG,CAC | GAA,TTC | GAT,ATC | GAC,GTC |
| GAA,TTC | GAC,GTC | CTT,AAG | CTT,AAG | GCG,CGC | CTT,AAG | GGG,CCC | GGC,GCC |
| GTG,CAC | GCG,CGC | GCG,CGC | GAA,TTC | ATG,CAT | GAT,ATC | GGA,TCC | GCG,CGC |
| GCG,CGC | GTG,CAC | GAA,TTC | GCG,CGC | GAT,ATC | GTG,CAC | GAG,CTC | GTG,CAC |
| ATG,CAT | GAT,ATC | GTG,CAC | GGT,ACC | GAA,TTC | ATG,CAT | CTT,AAG | CTT,AAG |
| GCA,TGC | CCA,TGG | GGT,ACC | GTG,CAC | CTT,AAG | GCG,CGC | GCT,AGC | GCT,AGC |
| GTA,TAC | GGT,ACC | GGG,CCC | GGG,CCC | GCT,AGC | GCT,AGC | GCA,TGC | GAA,TTC |
| CCA,TGG | GTA,TAC | CCG,CGG | CCG,CGG | GCA,TGC | CCA,TGG | ATG,CAT | GTA,TAC |
| GGT,ACC | GCA,TGC | GCA,TGC | CCA,TGG | GTA,TAC | GTA,TAC | GTA,TAC | GCA,TGC |
| GAT,ATC | ATG,CAT | CCA,TGG | GCA,TGC | GGT,ACC | GGT,ACC | GAA,TTC | ATG,CAT |
| GCT,AGC | GCT,AGC | CCT,AGG | CCT,AGG | CCA,TGG | GCA,TGC | CCG,CGG | CCG,CGG |
| CCG,CGG | CCG,CGG | TCG,CGA | TCG,CGA | CCG,CGG | CCG,CGG | CCT,AGG | CCT,AGG |
| CCT,AGG | CCT,AGG | GAG,CTC | GAG,CTC | CCT,AGG | CCT,AGG | CCA,TGG | CCA,TGG |
| TCG,CGA | TCG,CGA | TCT,AGA | TCT,AGA | TCG,CGA | TCG,CGA | TCG,CGA | TCG,CGA |
| TCA,TGA | TCA,TGA | ACG,CGT | ACG,CGT | TCA,TGA | TCA,TGA | TCA,TGA | TCA,TGA |
| TCT,AGA | TCT,AGA | ACT,AGT | ACT,AGT | TCT,AGA | TCT,AGA | TCT,AGA | TCT,AGA |
| ACG,CGT | ACG,CGT | ACA,TGT | ACA,TGT | ACG,CGT | ACG,CGT | ACG,CGT | ACG,CGT |
| ACT,AGT | ACT,AGT | ATG,CAT | GAT,ATC | ACT,AGT | ACT,AGT | ACT,AGT | ACT,AGT |
| ACA,TGT | ACA,TGT | GAT,ATC | ATG,CAT | ACA,TGT | ACA,TGT | ACA,TGT | ACA,TGT |
| CTG,CAG | CTG,CAG | CTG,CAG | CTG,CAG | CTG,CAG | CTG,CAG | CTG,CAG | CTG,CAG |
| CTA,TAG | CTA,TAG | GTA,TAC | GTA,TAC | CTA,TAG | CTA,TAG | CTA,TAG | CTA,TAG |
| GTT,AAC | GTT,AAC | CTA,TAG | CTA,TAG | GTT,AAC | GTT,AAC | GTT,AAC | GTT,AAC |
| CAA,TTG | CAA,TTG | CAA,TTG | CAA,TTG | CAA,TTG | CAA,TTG | CAA,TTG | CAA,TTG |
| ATA,TAT | ATA,TAT | ATA,TAT | ATA,TAT | ATA,TAT | ATA,TAT | ATA,TAT | ATA,TAT |
| ATT,AAT | ATT,AAT | ATT,AAT | ATT,AAT | ATT,AAT | ATT,AAT | ATT,AAT | ATT,AAT |
| TTA,TAA | TTA,TAA | TTA,TAA | TTA,TAA | TTA,TAA | TTA,TAA | TTA,TAA | TTA,TAA |
| TTT,AAA | TTT,AAA | TTT,AAA | TTT,AAA | TTT,AAA | TTT,AAA | TTT,AAA | TTT,AAA |

| 29 |
| --- |
| Li[42] |
| Direct codon chronology |
| Unambiguous |
| GGG,CCC |
| GGC,GCC |
| GGA,TCC |
| GAG,CTC |
| GAC,GTC |
| GGT,ACC |
| GCG,CGC |
| AGC,GCT |
| GCA,TGC |
| CGG,CCG |
| AGG,CCT |
| TGG,CCA |
| CGA,TCG |
| AGA,TCT |
| TGA,TCA |
| ACG,CGT |
| AGT,ACT |
| ACA,TGT |
| GTG,CAC |
| CAG,CTG |
| GAT,ATC |
| ATG,CAT |
| GAA,TTC |
| GTA,TAC |
| TAG,CTA |
| AAC,GTT |
| AAG,CTT |
| CAA,TTG |
| ATA,TAT |
| TAA,ATT |
| TAA,TTA |
| AAA,TTT |

#### SI-9: Strands surviving for reference codon chronologies

The codon chronologies calculated in SI-7 and shown in SI-8 lead to UV sensitivities of the associated proto-genome pools shown in SI-f1.

**SI-f1** Ratio of surviving strands for proto-genome pools plotted separately for all criteria used in Figure 4c. For codon chronologies that do not contain unambiguously sortable subsets, the limit plots for the chronologies compatible with the corresponding minimum and maximum UV sensitivity chronologies are shown using the same color.

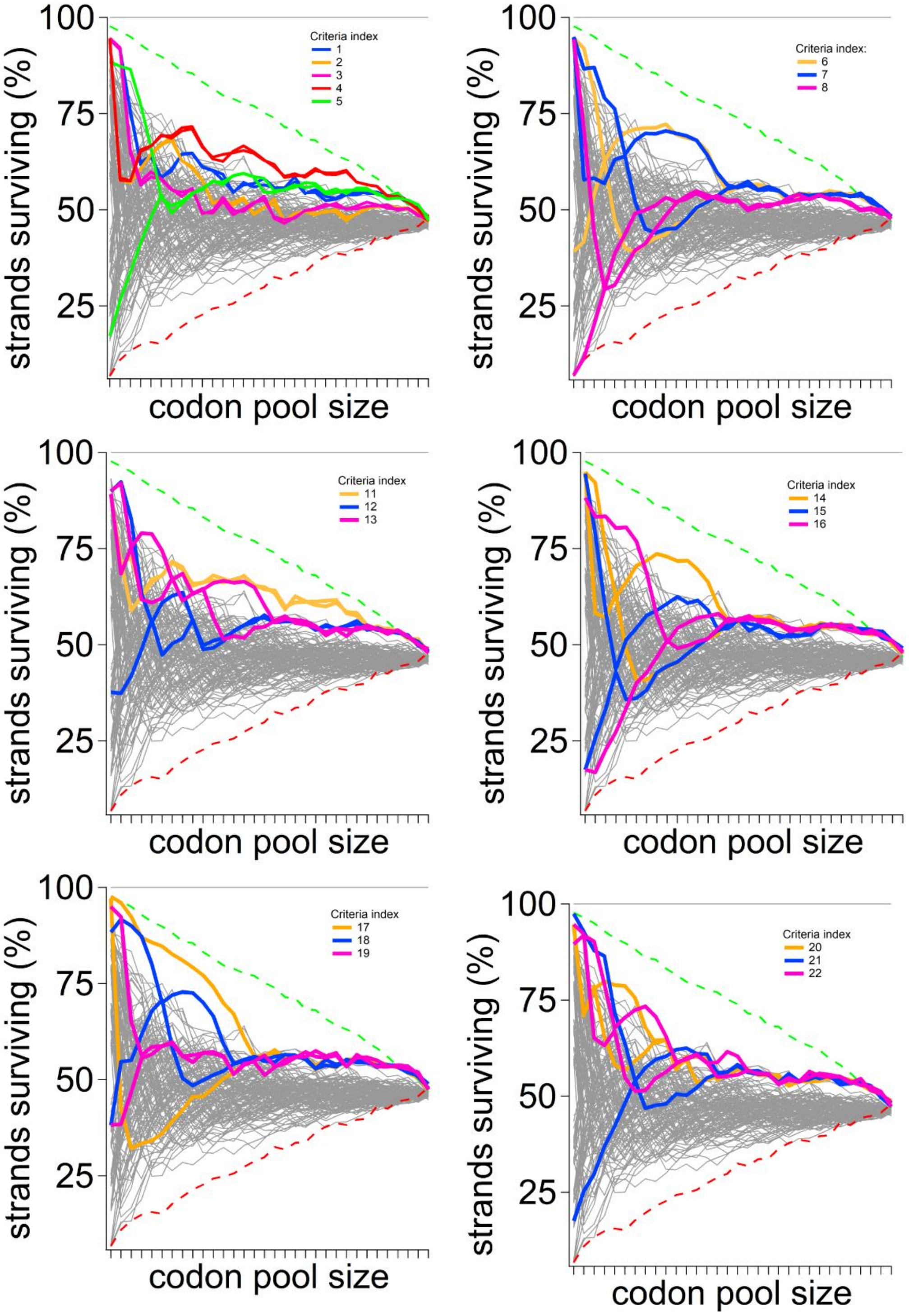

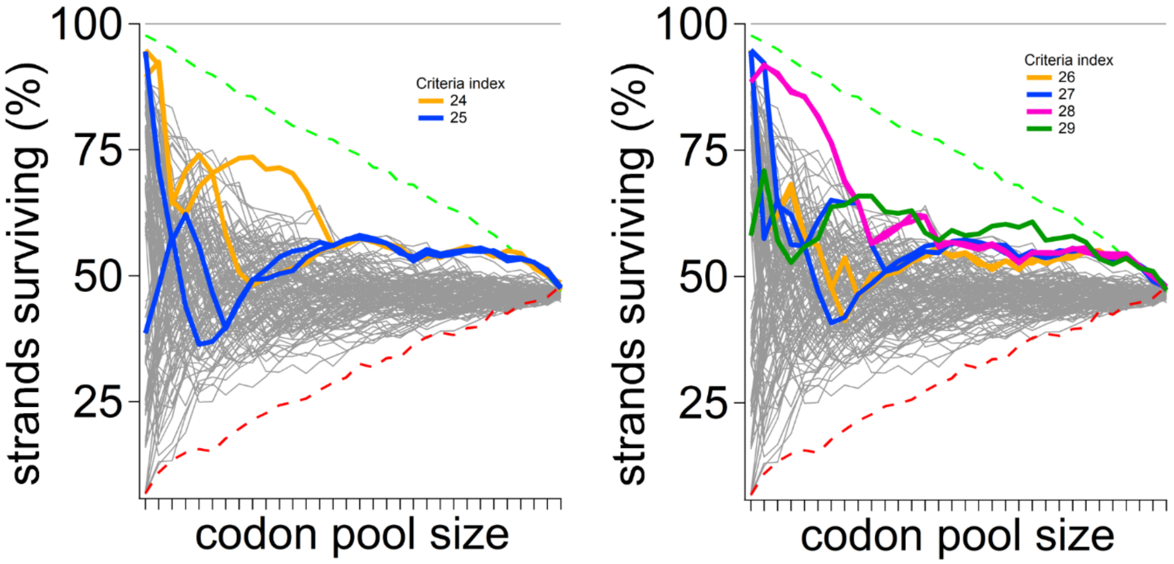


**References**


1 Lemaire, D. G. & Ruzsicska, B. P. Quantum yields and secondary photoreactions of the photoproducts of dTpdT, dTpdC and dTpdU. *Photochemistry and Photobiology* **57**, 755–769 (1993). https://doi.org:10.1111/j.1751-1097.1993.tb09207.x

2 Kumar, S. *et al.* Adenine photodimerization in deoxyadenylate sequences: elucidation of the mechanism through structural studies of a major d(ApA) photoproduct. *Nucleic Acids Res* **19**, 2841-2847 (1991). https://doi.org:10.1093/nar/19.11.2841

3 Law, Y. K., Azadi, J., Crespo-Hernández, C. E., Olmon, E. & Kohler, B. Predicting thymine dimerization yields from molecular dynamics simulations. *Biophysical Journal* **94**, 3590–3600 (2008). https://doi.org:10.1529/biophysj.107.118612

4 Sztumpf, E. & Shugar, D. Photochemistry of model oligo- and polynucleotides VI. Photodimerization and its reversal in thymine dinucleotide analogues. *Biochimica et Biophysica Acta (BBA) - Specialized Section on Nucleic Acids and Related Subjects* **61**, 555–566 (1962). https://doi.org:10.1016/0926-6550(62)90107-x

5 Johns, H. E., Pearson, M. L., LeBlanc, J. C. & Helleiner, C. W. The ultraviolet photochemistry of thymidylyl-(3′5′)-thymidine. *Journal of molecular biology* **9**, 503-IN501 (1964). https://doi.org:10.1016/s0022-2836(64)80223-0

6 Kaszubowski, J. D. & Trakselis, M. A. Beyond the Lesion: Back to High Fidelity DNA Synthesis. *Front Mol Biosci* **8**, 811540 (2021). https://doi.org:10.3389/fmolb.2021.811540

7 Kufner, C. L. *et al.* Sequence Dependent UV Damage of Complete Pools of Oligonucleotides. *bioRxiv*, 2022.2008.2001.502267 (2022). https://doi.org:10.1101/2022.08.01.502267

8 Tataurov, A. V., You, Y. & Owczarzy, R. Predicting ultraviolet spectrum of single stranded and double stranded deoxyribonucleic acids. *Biophys Chem* **133**, 66-70 (2008). https://doi.org:10.1016/j.bpc.2007.12.004

9 Pearce, B. K. D., Pudritz, R. E., Semenov, D. A. & Henning, T. K. Origin of the RNA world: The fate of nucleobases in warm little ponds. *Proc Natl Acad Sci U S A* **114**, 11327-11332 (2017). https://doi.org:10.1073/pnas.1710339114

10 Ranjan, S., Wordsworth, R. & Sasselov, D. D. Atmospheric Constraints on the Surface UV Environment of Mars at 3.9Ga Relevant to Prebiotic Chemistry. *Astrobiology* **17**, 687-708 (2017). https://doi.org:10.1089/ast.2016.1596

11 Schreier, W. J., Gilch, P. & Zinth, W. Early events of DNA photodamage. *Annual review of physical chemistry* **66**, 497–519 (2015). https://doi.org:10.1146/annurev-physchem-040214-121821

12 Banyasz, A. *et al.* Electronic excited states responsible for dimer formation upon UV absorption directly by thymine strands: joint experimental and theoretical study. *Journal of the American Chemical Society* **134**, 14834–14845 (2012). https://doi.org:10.1021/ja304069f

13 Trifonov, E. N. The triplet code from first principles. *Journal of Biomolecular Structure & Dynamics* **22**, 1-11 (2004).

14 Haig, D. & Hurst, L. D. A quantitative measure of error minimization in the genetic code. *J Mol Evol* **33**, 412-417 (1991). https://doi.org:10.1007/BF02103132

15 Tze-Fei Wong, J. Coevolution of genetic code and amino acid biosynthesis. *Trends in Biochemical Sciences* **6**, 33-36 (1981). https://doi.org:10.1016/0968-0004(81)90013-x

16 Papentin, F. On order and complexity. II. Application to chemical and biochemical structures. *Journal of Theoretical Biology* **95**, 225-245 (1982). https://doi.org:10.1016/0022-5193(82)90241-7

17 Dufton, M. J. Genetic code synonym quotas and amino acid complexity: cutting the cost of proteins? *J Theor Biol* **187**, 165-173 (1997). https://doi.org:10.1006/jtbi.1997.0443

18 Arques, D. G. & Michel, C. J. A complementary circular code in the protein coding genes. *J Theor Biol* **182**, 45-58 (1996). https://doi.org:10.1006/jtbi.1996.0142

19 Brooks, D. J., Fresco, J. R., Lesk, A. M. & Singh, M. Evolution of amino acid frequencies in proteins over deep time: inferred order of introduction of amino acids into the genetic code. *Mol Biol Evol* **19**, 1645-1655 (2002). https://doi.org:10.1093/oxfordjournals.molbev.a003988

20 Brooks, D. J. & Fresco, J. R. Increased frequency of cysteine, tyrosine, and phenylalanine residues since the last universal ancestor. *Mol Cell Proteomics* **1**, 125-131 (2002). https://doi.org:10.1074/mcp.m100001-mcp200

21 Eck, R. V. & Dayhoff, M. O. Evolution of the structure of ferredoxin based on living relics of primitive amino Acid sequences. *Science* **152**, 363-366 (1966). https://doi.org:10.1126/science.152.3720.363

22 Xia, T. *et al.* Thermodynamic parameters for an expanded nearest-neighbor model for formation of RNA duplexes with Watson-Crick base pairs. *Biochemistry* **37**, 14719-14735 (1998). https://doi.org:10.1021/bi9809425

23 Osawa, S., Jukes, T. H., Watanabe, K. & Muto, A. Recent evidence for evolution of the genetic code. *Microbiol Rev* **56**, 229-264 (1992). https://doi.org:10.1128/mr.56.1.229-264.1992

24 Crick, F. H., Brenner, S., Klug, A. & Pieczenik, G. A speculation on the origin of protein synthesis. *Orig Life* **7**, 389-397 (1976). https://doi.org:10.1007/BF00927934

25 Eigen, M. & Schuster, P. The Hypercycle. *The Science of Nature* **65**, 341-369 (1978). https://doi.org:10.1007/bf00439699

26 Eigen, M., Gardiner, W., Schuster, P. & Winkler-Oswatitsch, R. The origin of genetic information. *Sci Am* **244**, 88-92, 96, et passim (1981). https://doi.org:10.1038/scientificamerican0481-88

27 Ferreira, R. & Coutinho, K. R. Simulation studies of self-replicating oligoribotides, with a proposal for the transition to a peptide-assisted stage. *J Theor Biol* **164**, 291-305 (1993). https://doi.org:10.1006/jtbi.1993.1155

28 Nelsestuen, G. L. Amino acid-directed nucleic acid synthesis. A possible mechanism in the origin of life. *J Mol Evol* **11**, 109-120 (1978). https://doi.org:10.1007/BF01733887

29 Möller, W. & Janssen, G. M. C. Transfer RNAs for primordial amino acids contain remnants of a primitive code at position 3 to 5. *Biochimie* **72**, 361-368 (1990). https://doi.org:10.1016/0300-9084(90)90033-d

30 Chaley, M. B., Korotkov, E. V. & Phoenix, D. A. Relationships among isoacceptor tRNAs seems to support the coevolution theory of the origin of the genetic code. *J Mol Evol* **48**, 168-177 (1999). https://doi.org:10.1007/pl00006455

31 Henikoff, S. & Henikoff, J. G. Amino acid substitution matrices from protein blocks. *Proc Natl Acad Sci U S A* **89**, 10915-10919 (1992). https://doi.org:10.1073/pnas.89.22.10915

32 Eigen, M. & Winkler-Oswatitsch, R. Transfer-RNA, an early gene? *Naturwissenschaften* **68**, 282-292 (1981). https://doi.org:10.1007/BF01047470

33 Kvenvolden, K. A., Lawless, J. G. & Ponnamperuma, C. Nonprotein amino acids in the murchison meteorite. *Proc Natl Acad Sci U S A* **68**, 486-490 (1971). https://doi.org:10.1073/pnas.68.2.486

34 Riddle, D. S. *et al.* Functional rapidly folding proteins from simplified amino acid sequences. *Nat Struct Biol* **4**, 805-809 (1997). https://doi.org:10.1038/nsb1097-805

35 Yarus, M. Specificity of arginine binding by the Tetrahymena intron. *Biochemistry* **28**, 980-988 (1989). https://doi.org:10.1021/bi00429a010

36 Jukes, T. H. Possibilities for the evolution of the genetic code from a preceding form. *Nature* **246**, 22-26 (1973). https://doi.org:10.1038/246022a0

37 Wong, J. T. The evolution of a universal genetic code. *Proc Natl Acad Sci U S A* **73**, 2336-2340 (1976). https://doi.org:10.1073/pnas.73.7.2336

38 Hartman, H. Speculations on the origin of the genetic code. *J Mol Evol* **40**, 541-544 (1995). https://doi.org:10.1007/BF00166623

39 Taylor, F. J. R. & Coates, D. The code within the codons. *Biosystems* **22**, 177-187 (1989). https://doi.org:10.1016/0303-2647(89)90059-2

40 Ikehara, K., Omori, Y., Arai, R. & Hirose, A. A novel theory on the origin of the genetic code: a GNC-SNS hypothesis. *J Mol Evol* **54**, 530-538 (2002). https://doi.org:10.1007/s00239-001-0053-6

41 Baumann, U. & Oro, J. Three stages in the evolution of the genetic code. *Biosystems* **29**, 133-141 (1993). https://doi.org:10.1016/0303-2647(93)90089-u

42 Li, D. J. Formation of the Codon Degeneracy during Interdependent Development between Metabolism and Replication. *Genes (Basel)* **12** (2021). https://doi.org:10.3390/genes12122023
